# The theta paradox: 4-8 Hz EEG oscillations reflect both local sleep and cognitive control

**DOI:** 10.1101/2022.04.04.487061

**Authors:** Sophia Snipes, Elena Krugliakova, Elias Meier, Reto Huber

**Affiliations:** Child Development Centre, University Children’s Hospital Zürich, University of Zürich, Switzerland; Neural Control of Movement Lab, Department of Health Sciences and Technology, ETH Zürich; Department of Child and Adolescent Psychiatry and Psychotherapy, Psychiatric Hospital, University of Zürich, Switzerland

## Abstract

Human brain activity generates electroencephalographic (EEG) oscillations that characterize specific behavioral and vigilance states. The frequency of these oscillations is typically sufficient to distinguish a given state, however theta oscillations (4-8 Hz) have instead been found in near-opposite conditions of drowsiness during sleep deprivation and alert cognitive control. While the latter has been extensively studied and is often referred to as “frontal midline theta”, the former has been investigated far less but is considered to be a marker for local sleep during wake. In this study we investigated to what extent theta oscillations differed during cognitive tasks and sleep deprivation. We measured high-density EEG in 18 young healthy adults performing 6 tasks under 3 levels of sleep deprivation. We found both cognitive load and sleep deprivation increased theta power in medial prefrontal cortical areas, however sleep deprivation caused additional increases in theta in many other, predominantly frontal, areas. The sources of sleep deprivation theta were task-dependent, with a visual-spatial task and short-term memory task showing the most widespread effects. Notably, theta was highest in supplementary motor areas during passive music listening, and highest in the inferior temporal cortex during a spatial game. This suggests that theta caused by sleep deprivation may preferentially occur in cortical areas not involved in ongoing behavior. While our results find differences in topography from frontal midline theta, they raise the possibility that a common mechanism may underly both theta oscillations during cognition and during sleep deprivation.

## INTRODUCTION

Research of the healthy human brain is limited to what can be measured non-invasively. Electroencephalography (EEG) offers the possibility of measuring synchronized neural activity, i.e. oscillations. Since the discovery of EEG, specific oscillations have been associated with behavioral states such as alertness, drowsiness, and different stages of sleep. This allows oscillations to be used as objective markers for vigilance. While the exact function of a given oscillation is not always understood, its predominant occurrence is limited to a specific vigilance state. The exception are theta oscillations (4-8 Hz), which have been separately identified as an indicator of drowsiness and intense cognition.

Theta oscillations are associated with drowsiness in both animals (Vyazovskiy & Tobler, 2005) and humans (Finelli et al., 2000), most easily observed during sleep deprivation. More specifically, theta activity reflects sleep pressure, i.e. the interaction between circadian rhythm and time spent awake that determines when an individual feels the need to sleep (Aeschbach et al., 1997; Borbely, 1982; Cajochen et al., 2002; Finelli et al., 2000; Strijkstra et al., 2003). Furthermore, theta activity is elevated in patients with excessive daytime sleepiness (Grenèche et al., 2008).

Not only does theta track sleep pressure in time, but also in space. Sleep pressure appears driven by daytime neural plasticity; brain areas that undergo high plasticity during the day will have higher sleep need at the beginning of the night, reflected as a local increase in slow wave activity (1-4 Hz oscillations) during deep sleep (Tononi & Cirelli, 2014). This increased sleep pressure is especially true for frontal association areas, where both theta and slow waves are highest (Finelli et al., 2000). Similarly, daytime activity can induce local differences in sleep pressure in areas involved in learning, leading to corresponding local changes in slow wave activity (Huber et al., 2004, 2006). The tight relationship between daytime activity and local changes in theta was convincingly illustrated in a study by Hung et al. (2013), in which participants spent a 24 h sleep deprivation period either playing a driving simulator or listening to audiobooks. Theta during waking rest between tasks increased more over visual (occipital/parietal) areas after playing the driving simulator, and more over auditory/speech (left temporal) areas after listening to audiobooks.

Given the presence of theta oscillations when and where sleep pressure is highest, they have been hypothesized to be a form of local sleep during wake (Bernardi & Siclari, 2019; Siclari & Tononi, 2017; Vyazovskiy et al., 2011), specifically localized slow waves. During sleep, slow waves in the surface EEG correspond to synchronized silencing of neuronal spiking in large populations of neurons, known as “off periods” (Steriade et al., 2001). Vyazovskiy et al. (2011) found these off periods to also occur during sleep deprived awake rats, over localized cortical areas. These off periods in wake corresponded to theta oscillations in the local field potentials. While evidence for sleep slow wave off periods in humans has been found with intracortical recordings in epileptic patients (Nir et al., 2011), the same has not been possible for theta in wake due to technical limitations in single-cell recordings in humans (Nir et al., 2017). It is therefore still uncertain if theta in humans corresponds to off periods in the cortex and could thus be considered localized slow waves.

Equally robust research has separately linked theta activity to cognition and wake. Theta has been associated with an unparalleled variety of functions (for a review see Buzsáki, 2005), most notably hippocampal theta during spatial navigation in rats (Buzsáki, 1996; O’Keefe & Recce, 1993) and frontal-midline theta during cognitive tasks in humans. Frontal-midline theta (fmTheta) is an oscillation that is visually identifiable in approximately 10-40% of the population (Inanaga, 1998), occurs in bursts around 1-10 seconds, with amplitudes around 30-60 μV (Mitchell et al., 2008). It has been associated with arithmetic (Ishihara & Yoshii, 1967; Ishii et al., 2014), working memory (Gevins et al., 1998; Jensen & Tesche, 2002), and even meditation (Banquet, 1973; D. J. Lee et al., 2018). It has been shown to increase with short-term / working memory load. fmTheta has been source localized to the anterior cingulate cortex and medial prefrontal cortex (Ishii et al., 2014; Michels et al., 2010; Onton et al., 2005), where it has in turn been anti-correlated to fMRI BOLD (functional magnetic resonance imaging, blood-oxygen level dependent) activity in these areas (Scheeringa et al., 2008, 2009).

The exact function of fmTheta oscillations in cognition is still unresolved although various explanations have been proposed (Anderson & Hulbert, 2021; Bastiaansen & Hagoort, 2003; Hsieh & Ranganath, 2014; Klimesch et al., 2005; Lisman & Jensen, 2013; Raghavachari et al., 2001; Sauseng et al., 2010). One of the most well-elaborated hypotheses is that theta is responsible for synchronizing neuronal firing to specific phases of distant cortical regions (Lisman & Jensen, 2013). Such a role for theta has been supported by intracortical recordings in macaques for short term memory tasks (H. Lee et al., 2005; Liebe et al., 2012), however intracortical recordings in humans have produced mixed results, with multiple areas showing independent task-related changes in theta (Brzezicka et al., 2018). Nevertheless, given the strong association between theta and tasks, it is generally considered to be functionally relevant for cognitive processing.

Currently, research in theta oscillations caused by sleep deprivation (hereafter referred to as sdTheta) has remained largely independent from research in cognition and fmTheta. sdTheta has been studied almost exclusively during quiet resting states, whereas fmTheta is studied during cognitive tasks under low sleep pressure conditions (i.e. normal wake). It is therefore unknown if these represent either two distinct oscillations in the theta range or one and the same, as has been suggested by Takahashi et al. (1997) and Mitchell et al. (2008). If sdTheta and fmTheta are qualitatively distinct, this would resolve the apparent paradox of an oscillation reflecting both drowsiness and cognition. If sdTheta is instead a manifestation of fmTheta, then its interpretation as local sleep should be reconsidered. Instead, fmTheta during sleep deprivation would speak of some form of compensation mechanism counteracting the detrimental effects of prolonged wakefulness.

We conducted a sleep deprivation study in young healthy adults to disentangle the changes in theta related to both drowsiness and cognition using high-density EEG. Participants performed 6 tasks at 3 time points: a baseline session (**BL**) in the morning after a normal night of sleep, a sleep restriction session (**SR**) conducted at the same time of day as BL but after only 4 hours of sleep, and a sleep deprivation session (**SD**) session after 20 h of wake following the 4 h night of sleep.

The tasks included:

- A short-term memory task (**STM**) to reproduce classic fmTheta memory load effects
- The psychomotor vigilance task (**PVT**) which involved responding to the start of a countdown every 2-10 s with the push of a button
- The lateralized attention task (**LAT**), an adaptation of the PVT in which faint circles would appear in half of the screen to which participants had to push a button in response
- A speech fluency task (**Speech**) in which participants had to read out loud English tongue twisters
- The Arkanoid-based game BBTAN (**Game**) which involved aiming a ball to bounce off bricks to make them disappear before they covered the screen
- Passive music listening (**Music**)

To determine whether sdTheta and fmTheta could be considered the same oscillation, we first looked at their topography within the STM task, and with source localization identified their neural substrates. We then investigated how they interact by reproducing the classic fmTheta memory load effect in the STM task at each session. Then, to determine more generally how sdTheta is affected by behavioral state, we compared its topography and source localization in all 6 tasks. In a final attempt to distinguish fmTheta from sdTheta, we inspected the spectrograms for each task to determine if they could be differentiated by peak frequency. Altogether, these results provide the first look of theta simultaneously under different behavioral and vigilance states.

## RESULTS

### Changes in sleep architecture and subjective sleepiness confirm the effectiveness of the sleep deprivation protocol

To determine whether the sleep deprivation protocol was successful in increasing sleep pressure, we evaluated changes in sleep architecture between the baseline night and recovery night following sleep deprivation (Table 1). We found shorter sleep onset latencies and more deep sleep (NREM3), key indicators of increased sleep pressure.

**Table 1:** Sleep architecture. All values in the first three columns are in mean minutes ± standard deviations, except SE which is in percentages (100 & SDU/ Total time in bed). The last three columns indicate p-values from paired t-tests (α=5%) between the different nights. Acronyms: REM (rapid eye movements), SOL (sleep onset latency), SDU (sleep duration), WASO (wake after sleep onset), SE (sleep efficiency), ROL (REM onset latency), NIGHTPRE (night before sleep deprivation period), NIGHTPOST (night after sleep deprivation period).

|  | BASELINE | NIGHTPRE | NIGHTPOST | NIGHTPRE<br>VS BASELINE | NIGHTPOST<br>VS<br>BASELINE | NIGHTPOST<br>VS<br>NIGHTPRE |
| --- | --- | --- | --- | --- | --- | --- |
| WAKE | 40.6 ± 30.1 | 18.1 ± 17.2 | 20.7 ± 9.3 | .001 | .008 | .544 |
| NREM1 | 22.3 ± 12.3 | 10.8 ± 8.8 | 10.6 ± 4.8 | < .001 | < .001 | .919 |
| NREM2 | 257.6 ± 34.2 | 115.1 ± 22.3 | 214.3 ± 51.6 | < .001 | .001 | < .001 |
| NREM3 | 89.7 ± 40.1 | 64.6 ± 23.8 | 109.6 ± 37.0 | < .001 | .027 | < .001 |
| REM | 111.8 ± 32.0 | 33.4 ± 13.2 | 118.2 ± 29.8 | < .001 | .450 | < .001 |
| SOL | 16.8 ± 7.8 | 16.9 ± 13.2 | 5.6 ± 2.1 | .986 | < .001 | .002 |
| SDU | 481.8 ± 31.1 | 224.2 ± 16.8 | 453.0 ± 76.2 | < .001 | .108 | < .001 |
| WASO | 28.1 ± 26.0 | 5.7 ± 8.4 | 17.1 ± 9.0 | .001 | .076 | .001 |
| SE (%) | 92.5 ± 5.2 | 92.6 ± 7.0 | 95.6 ± 1.8 | .950 | .018 | .080 |
| ROL | 103.5 ± 47.5 | 98.7 ± 41.2 | 60.9 ± 19.4 | .458 | < .001 | < .001 |

All sleep stages except REM sleep showed a significant change between baseline and recovery, with NREM3 increasing 30% at the expense of wake (−30%), NREM1 (−47%), and NREM2 (−16%). Sleep onset latency (SOL) significantly decreased from 16.8 minutes to 5.6 minutes. Overall sleep duration was shorter during the recovery night, although this was not statistically significant (p-value = .108), and sleep efficiency increased from 92% to 96%. Together, these results indicate that sleep pressure, specifically for slow wave sleep, increased over the 24h wake period.

To determine the degree of sleep deprivation experienced by the participants, a brief questionnaire was administered after each task at every session (all questionnaire results are provided in Supplementary Figure 1). Participants indicated on a continuous Karolinska Sleepiness Scale (Åkerstedt & Gillberg, 1990) how alert or sleepy they felt (Figure 1A). A twoway repeated measures analysis of variance (rmANOVA) was conducted with factors *session* and *task*, as well as their interaction. There was a highly significant and extremely large effect of session (F_(2, 30)_ = 35.42, p < .000, η^2^ = .355), a significant medium effect of task (F_(5, 75)_ = 14.7, p < .000, η^2^ = .073) and a non-significant interaction (F_(10, 150)_ = 0.96, p = .440, η^2^ = .008). With sleep deprivation, participants felt less sleepy during the Game and most sleepy during the STM task (Figure 1B).

**Figure 1:**
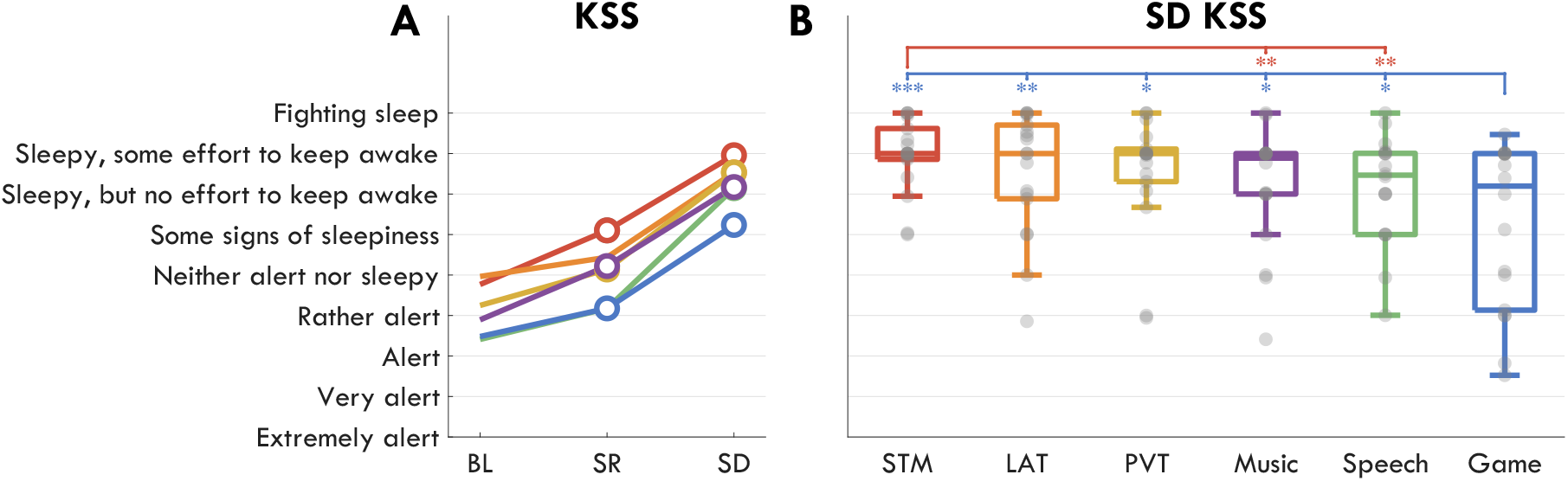
Subjective sleepiness ratings on the KSS following each task during each session. **A**: Average scores. Each colored line represents a task (STM: red, LAT: orange, PVT: yellow, Speech: green, Game: blue, Music: purple). Filled circles indicate a significant change from BL. **B**: KSS scores during SD. Gray circles represent each participant, the boxplot indicates median and interquartile range. Stars indicate significant differences, FDR corrected for multiple comparisons, between tasks (the color indicates one task, the location of the stars the other) such that: * p-value < .05, ** p-value < .01, *** p-value < .001. The empty tick mark indicates a trend p-value < .1). Acronyms: KSS (Karolinska Sleepiness Scale), BL (baseline), SR (sleep restriction), SD (sleep deprivation), STM (short term memory task), LAT (lateralized attention task), PVT (psychomotor vigilance task).

### Frontal-midline theta is more localized than sleep deprivation theta

For fmTheta and sdTheta to be considered the same oscillation, they should originate from the same brain areas. To determine if this was the case, we analyzed changes in theta from the short-term memory task (STM). Briefly, participants were presented with 1, 3, or 6 symbols for 2 s, which they then had to hold in memory during a 4 s retention period. Finally, they were presented a probe symbol and had to indicate whether it was part of the original set or not. Each session consisted of 120 randomized trials, 40 for each memory load level (**L1, L3, L6**).

Theta activity was measured as power spectral density (PSD) from the first 2 s of the retention period. To account for large interindividual differences (Suppl. Figure 2) as well as the 1/f power amplitude distribution across frequencies, PSD values were z-scored for each frequency, then the PSD for frequencies between 4 and 8 Hz was averaged. Power spectra for the retention period across load levels and sessions is provided in Suppl. Figure 3.

Source localization was performed with dynamic imaging of coherent sources (DICS) beamformers approach (Gross et al., 2001; Westner et al., 2022). After projecting untransformed theta power to the source space, trials were averaged by load level for each session and the data was z-scored across voxels, load, and sessions. Separate pipelines (Suppl. Figure 12) were then conducted for the visualization of approximate sources on an inflated brain (Figure 2 II-V), and for the labelling of anatomical sources (Figure 2C).

**Figure 2:**
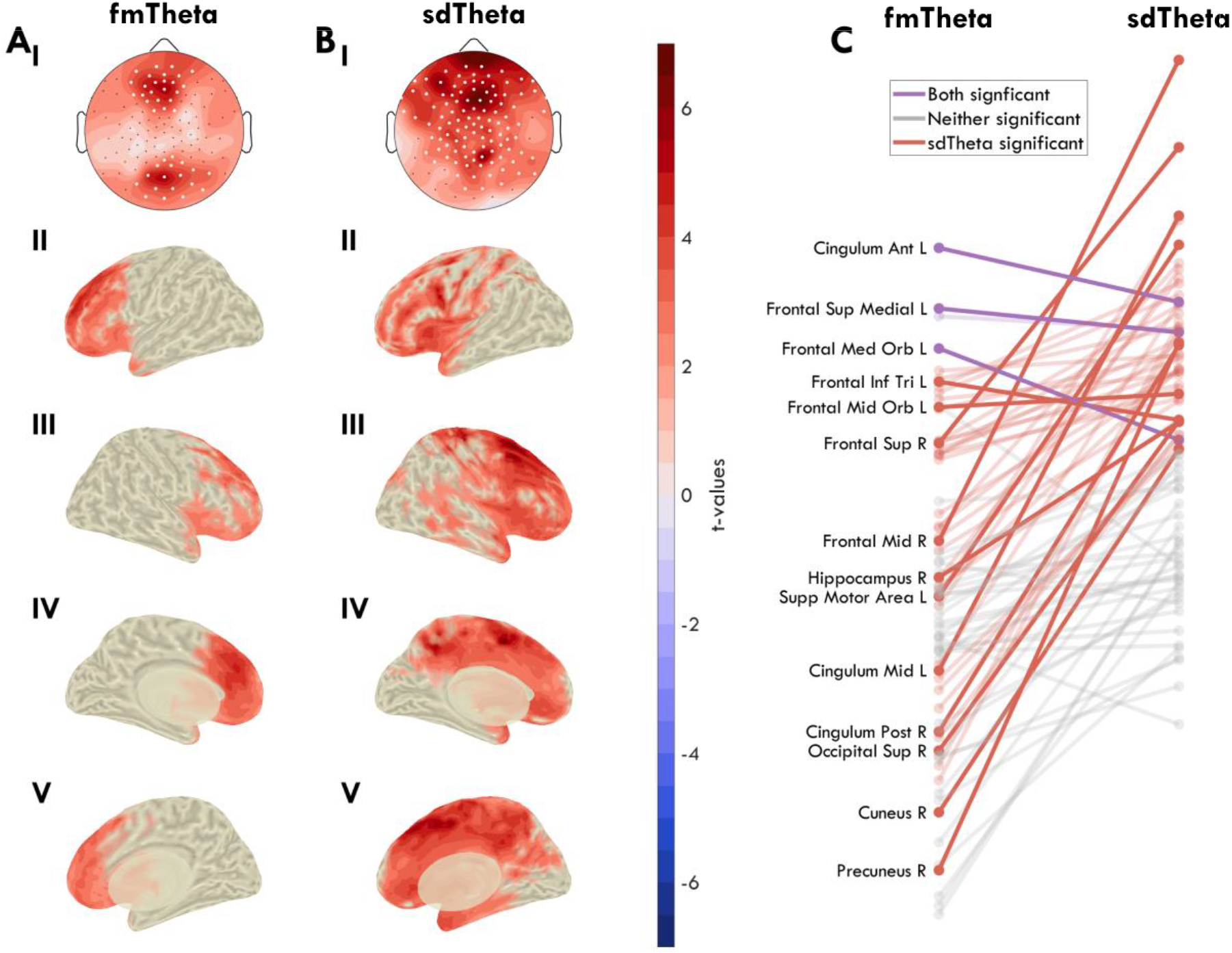
Average z-scored theta power (4-8 Hz) during the first 2 s retention window of the STM task. **A**: Frontal-midline theta, calculated as the difference between trials with 3 items vs 1 item to hold in memory, from the BL session. **B**: Sleep deprivation theta, calculated as the difference between SD trials and BL trials with 1 item to hold in memory. **I**: Theta changes represented in a 2D topography of EEG channels, as a head seen from above. Black dots indicate all channels, white dots indicate channels in which the change was statistically significant (p-value < .05) based on paired t-tests, FDR corrected for multiple comparisons. Source localization is presented in II-V as inflated brains. T-values are plotted with the same color scale in the channel and source space, such that red indicates a positive increase in theta from L1 to L3 in A, and from BL to SD in B. In the source space, voxel-wise cluster correction was implemented to mask non-significant effects. **II**: Left hemisphere, lateral view. **III**: Right hemisphere, lateral view. **IV**: Left hemisphere, medial view. **V**: Right hemisphere, medial view. **C**: Change in t-values for all areas between the fmTheta (A) and the sdTheta (B) comparisons, based on the AAL atlas. Lines in gray depict areas that showed no significant effects in either comparison, after FDR correction. Lines in red indicate areas showing a significant change in sdTheta, and lines in purple both in sdTheta and fmTheta. No area was only significant for fmTheta. Exact t-values can be seen in Figure 6. Acronyms: BL (baseline), SD (sleep deprivation), FDR (false discovery rate), STM (short term memory task), AAL (Automated Anatomical Labelling).

fmTheta was calculated comparing L3 trials to L1 trials for every channel at BL (Figure 2A). In the channel space, two significant channel groups emerged: the frontal peaking over ch11 (Fz) with t = 5.61, p = .002, Hedge’s g = 0.76; the posterior peaking over ch75 (Oz) with t = 5.61, p = .002, g = 0.76. Source localization identified the left medial frontal cortex as the main source (Figure 2A IV), especially the anterior cingulate cortex (t = 4.76) and the superior frontal gyrus, medial (t = 4.06) as well as orbital part (t = 3.59; all t-values provided in Figure 6). The right medial cortex also showed increases in theta however these areas did not survive correction for multiple comparisons.

sdTheta was calculated comparing L1 trials from BL to L1 trials from SD (Figure 2B). sdTheta was more widespread across the cortex than fmTheta, showing cluster-corrected increases in 38% of gray matter voxels relative to 21%, respectively. All areas showing load-effects of fmTheta were also significant for sdTheta (Figure 2C), and the areas showing highest sdTheta were not among those significantly increasing in fmTheta. Specifically, the peak location of sdTheta was different in both the channel space (ch5) and source space: right middle frontal gyrus (t = 6.95) and superior frontal gyrus (t = 5.94; 2B III). sdTheta extended along the medial cortex up to the cuneus (t_max_^1^ = 5.15) and was additionally present around the left insula (t_max_ = 4.58), and the temporal poles (t_max_ = 3.67). Therefore, sdTheta and fmTheta have different primary sources, and different spread throughout the cortex.

### Frontal-midline theta fades with increasing sleep deprivation

If sdTheta and fmTheta are independent oscillations, they should both be present during sleep deprivation when performing the STM task. fmTheta was therefore calculated at every session, for both L3 vs L1 and L6 vs L1 (Figure 3A). Surprisingly, fmTheta decreased in amplitude with increasing sleep deprivation, until no channel showed statistically significant differences with memory load during SD.

**Figure 3:**
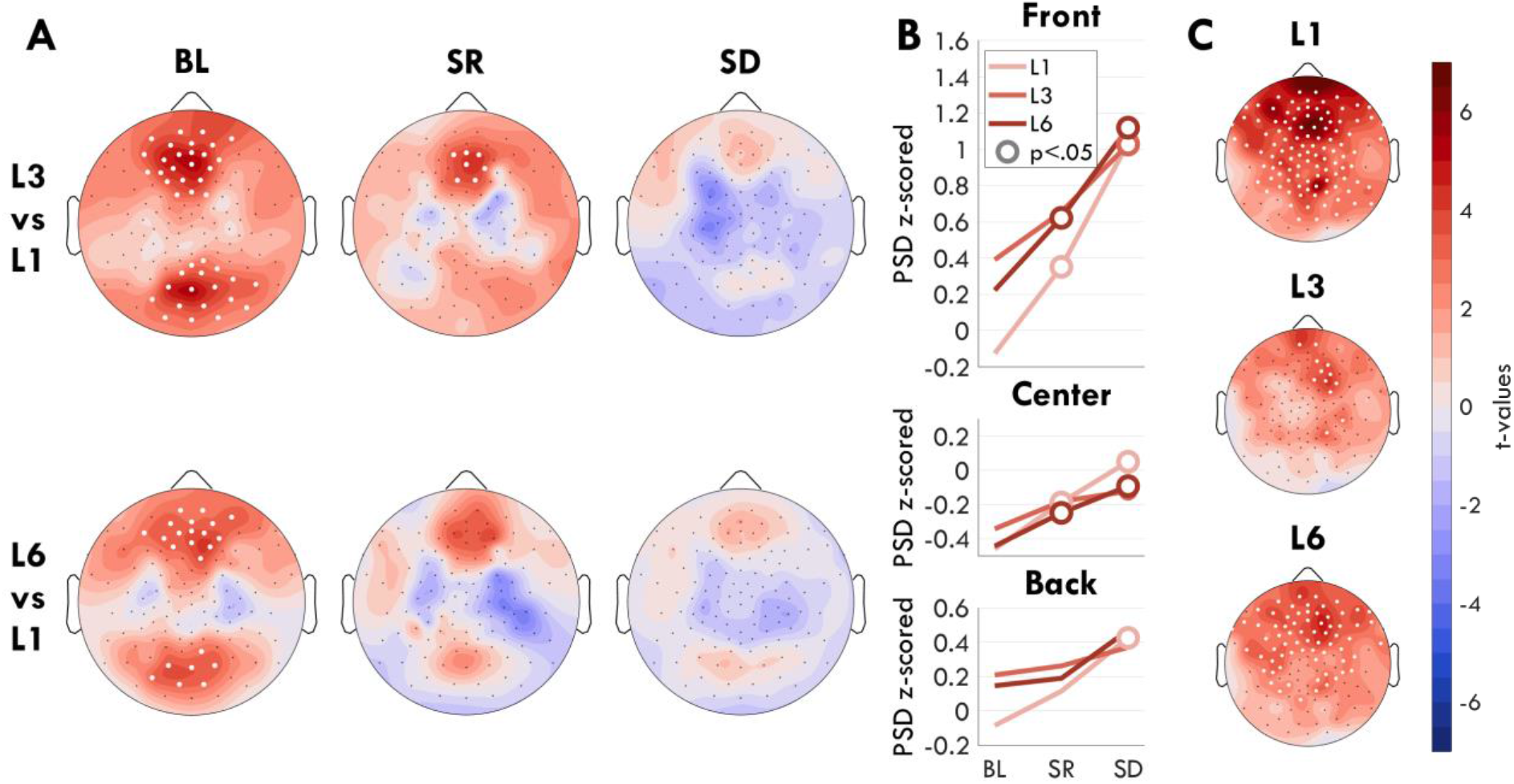
**A**: Difference in theta power for the first half of the retention period (2 s) during the STM task between level 3 (top row) and level 6 (bottom row) relative to level 1 for every session. Color represents t-values such that red indicates greater theta power in L3/L6 relative to L1 (same color scale as C). White dots indicate a significant effect, FDR corrected for multiple comparisons. **B**: Mean z-scored theta power across sessions for each load level. Each plot reflects a separate ROI. White circles indicate a significant change from BL, filled circles a trend, FDR corrected for multiple comparisons. **C**: Change in theta power during the retention periods between SD and BL, split by memory load. Acronyms: PSD (power spectral density), STM (short term memory task), ROI (region of interest), BL (baseline), SR (sleep restriction), SD (sleep deprivation), FDR (false discovery rate).

A two-way rmANOVA was conducted with factors *session, load*, and their interaction, separately for three regions of interest (ROIs): front, center, and back channels. In the front ROI there was both a significant and large effect of session (F_(2, 34)_ = 17.17, p < .001, eta2 = .287), a significant but small effect of load (F_(2, 34)_ = 5.92, p = .008, eta2 = .030), and a small significant interaction (F_(4, 68)_ = 3.74, p = .017, eta2 = .016). In the center ROI there was a significant effect of session (F_(2, 34)_ = 10.16, p < .001, eta2 = 0.198), no effect of load (F_(2, 34)_ = 1.35, p = .271, eta2 = .006), and a trending interaction (F_(4, 68)_ = 2.37, p = .095, eta2 = .022). In the back ROI there was a significant effect of session (F_(2, 34)_ = 4.64, p = .028, eta2 = .072), a small trending effect of load (F_(2, 34)_ = 2.63, p = .096, eta2 = 0.013), and a significant interaction (F_(4, 68)_ = 3.88, p = .019, eta2 = .014).

The interaction between load and session was driven by a larger increase in theta for low memory load trials during sleep deprivation (Figure 3B). To better understand this, we compared sdTheta topographies (BL vs SD) for each memory load level (Figure 3C). L1 showed the largest and most widespread increase in theta (t_max_ = 7.28, p < .001, g = 1.57), L3 the lowest and most local increase (t_max_ = 4.32, p = .024, g = 0.87), and L6 was intermediate (t_max_ = 4.93, p = .007, g = 1.45). As a result of sdTheta increasing more in low memory load trials, fmTheta effectively disappeared.

### Sources of sdTheta are task dependent

The results from Figure 2 show distinct topographies for fmTheta and sdTheta. The literature has identified fmTheta to consistently originate from the same medial region, however similar source localization has never been done for sdTheta. To determine whether the location of sdTheta is consistent or task-dependent we compared theta changes from BL in 6 different tasks, using the first 4 min of EEG data from each task. Figure 4 depicts the sdTheta changes for both SR and SD relative to BL in the channel space, and Figure 5 provides the source localization for SD relative to BL. Figure 6 provides the t-values for all regions found to be significant in at least one comparison of SD relative to BL. Mean theta values for all tasks in regions of interest are provided in the supplementary material (Suppl. Figure 4), as well as statistical comparisons between tasks (Suppl. Table 2).

**Figure 4:**
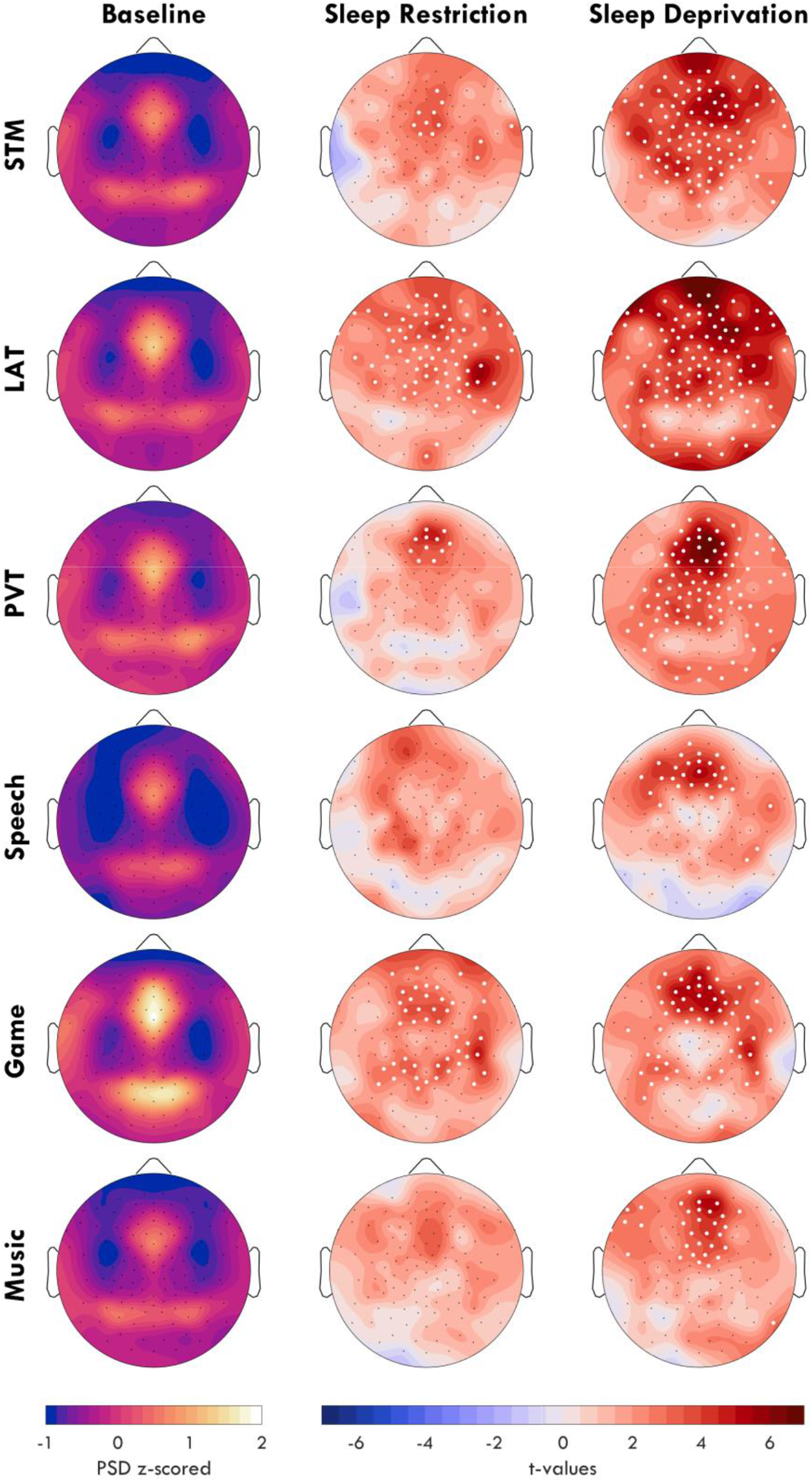
**First column**: mean z-scored theta power topographies at BL. **Second & third column**: the change in theta power from BL to SR and SD. Color indicates t-values, with red indicating an increase relative to BL, and blue a decrease. Black dots indicate all channels, white dots indicate channels in which the change was statistically significant (p-value < .05), FDR corrected for multiple comparisons. Acronyms: BL (baseline), SR (sleep restriction), SD (sleep deprivation), PSD (power spectral density), STM (short term memory task), LAT(Lateralized attention task), PVT (psychomotor vigilance task), FDR (false discovery rate).

**Figure 5:**
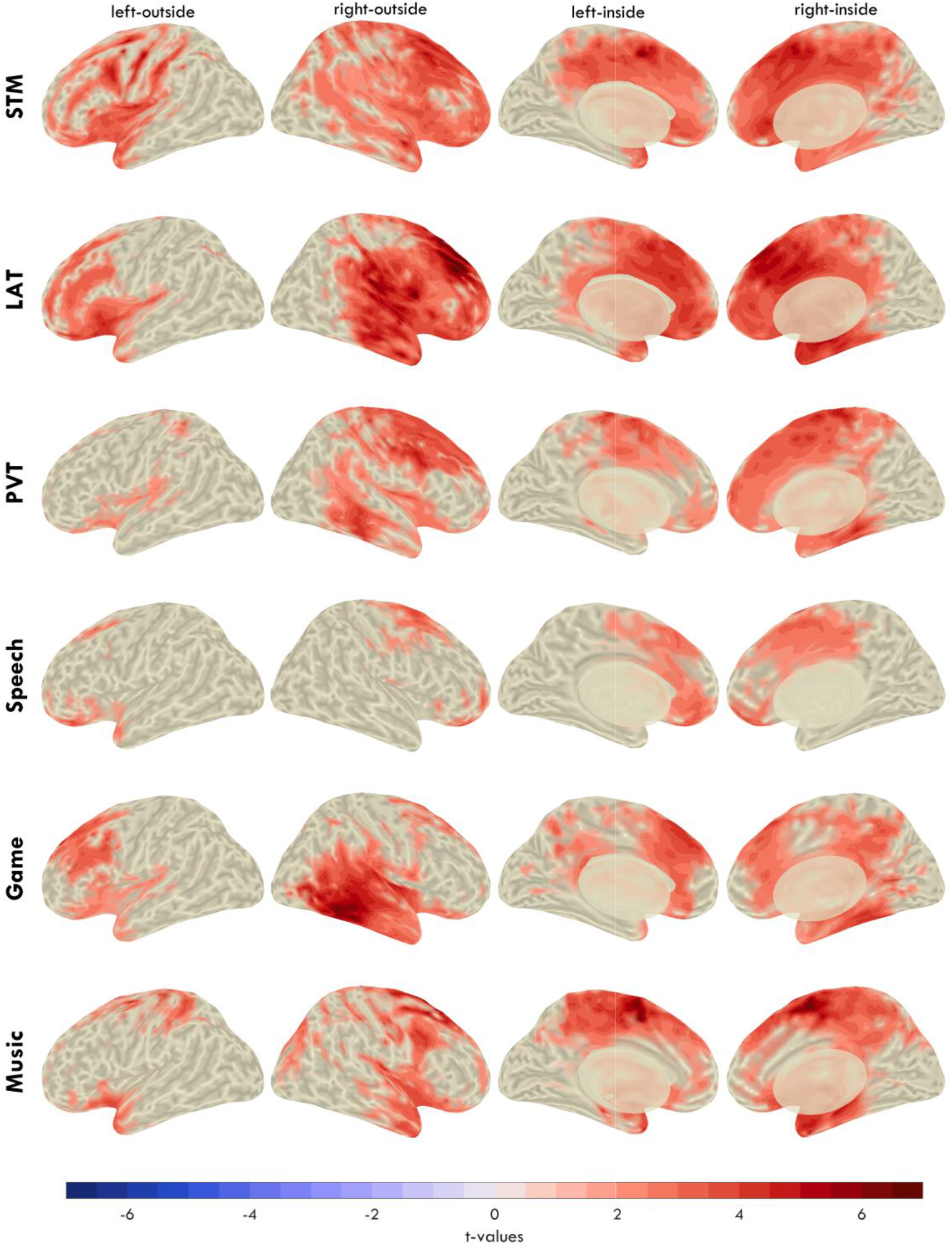
Change in theta from BL to SD from Figure 4 in the source space, projected onto the inflated brain. Voxel-wise cluster correction was implemented to mask non-significant effects. Color indicates t-values, such that red indicates an increase in power from BL to SD. Acronyms: BL (baseline), SD (sleep deprivation), STM (short term memory task), LAT (Lateralized attention task), PVT (psychomotor vigilance task).

**Figure 6:**
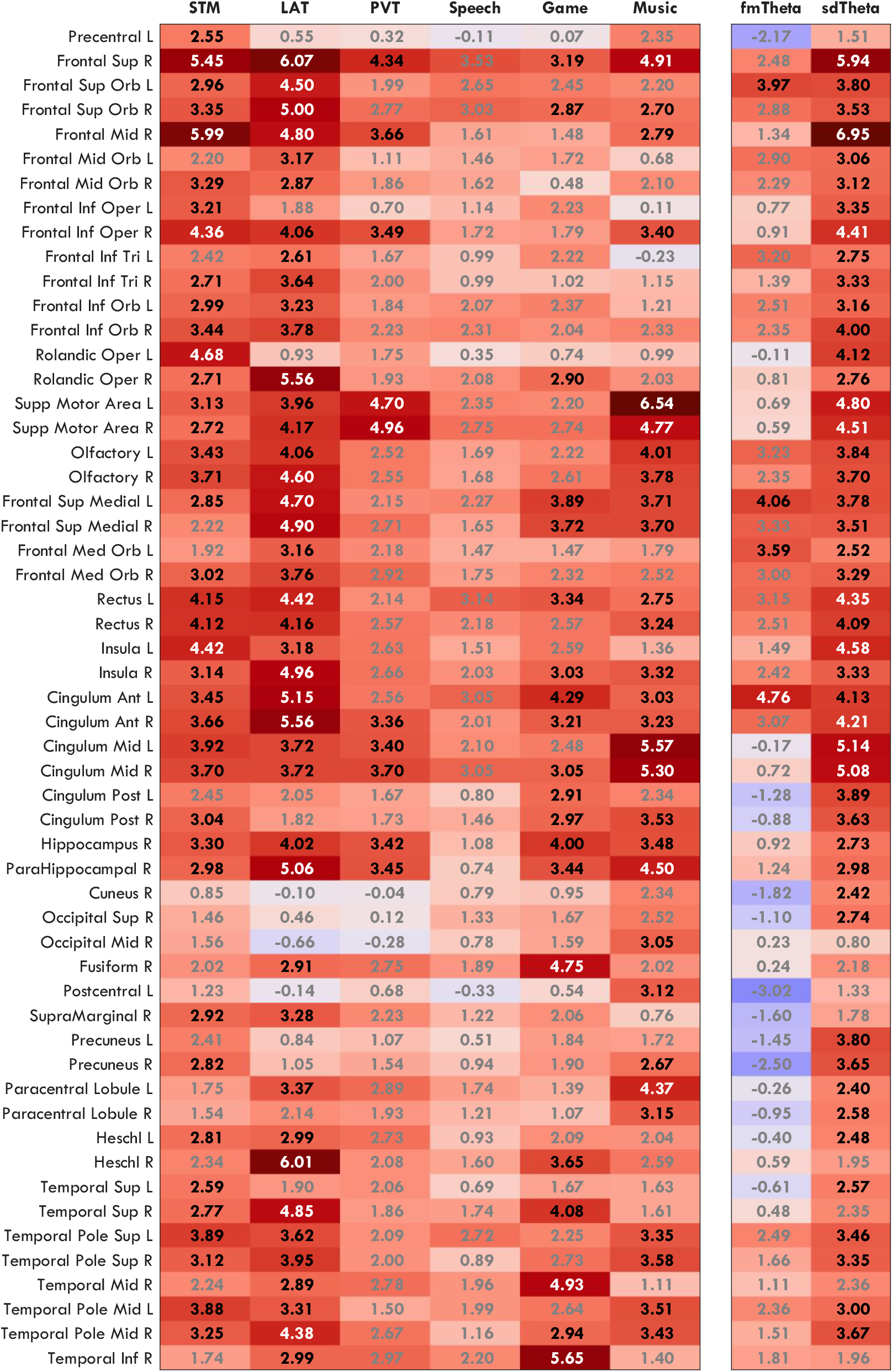
T-values for all areas from the AAL atlas that showed a significant change in at least one comparison (BL vs SD in all tasks; L3 vs L1 from fmTheta and L1 BL vs L1 SD for sdTheta in the STM). Text in gray indicates areas not significant after FDR correcting for multiple comparisons. Text in white indicates the top 10% t-values. Acronyms: AAL (Automated Anatomical Labelling), BL (baseline), SD (sleep deprivation), STM (short term memory task), LAT (Lateralized attention task), PVT (psychomotor vigilance task).

All tasks showed increases in theta between BL and SR, however no channel was significant for the Speech and Music conditions after FDR correction. The highest overall increase was seen for the LAT over ch109 (t_max_ = 5.74, p = 0.002, g = 1.33), accompanied by widespread increases. Due to the otherwise medium-low effect sizes, the comparison between BL and SR was not further investigated with source localization.

For BL to SD, already from the channel space it is evident that the location and spread of sdTheta is task dependent. The LAT, STM, and PVT showed the most widespread increases, as well as the highest amplitude (PVT: t_max_ = 7.52, p < .001, g = 1.85; LAT: t_max_ = 7.10, p < .001, g = 1.24; STM: t_max_ = 6.31, p = .001, g = 1.80). The Speech task showed the lowest and most local increase in theta (t_max_ = 5.50, p = .005, g = 1.51).

The source space allowed further anatomical localization of the origin of theta activity. All tasks showed a predominantly right, frontal increase in theta, although no anatomical area survived FDR correction for the Speech task. One of the primary sources of sdTheta across all tasks was the right superior frontal gyrus. All tasks (except Speech) also had significant theta originating from the right hippocampus, parahippocampus, anterior and middle cingulate cortex. The STM and LAT had further extensive increases across both dorsal and medial frontal areas, with the STM showing high theta activity along the left lateral sulcus (Rolandic operculum, insula), and the LAT in the right lateral sulcus (Heschl’s gyrus, Rolandic operculum, insula). Unfortunately, source localization along this sulcus is challenging due to how gray matter is folded and would require subject-specific MRI structural scans for accurate results.

The overall strongest source of sdTheta was the left supplementary motor area during the Music task (t = 6.54), extending contralaterally as well as into the middle cingulate cortex. Bilateral supplementary motor areas were also the main sources of theta for the PVT (t_max_ = 4.96). The supplementary motor area showed significant increases in the LAT and STM but to a lesser extent (STM: t_max_ = 3.66; LAT: t_max_ = 4.17) and were not significant in the Game (t_max_ = 2.74).

Finally, the most atypical distribution of sdTheta came from the Game, which showed minimal increases in frontal cortices and primary sdTheta originating from the right inferior temporal cortex (inferior temporal gyrus, mid temporal gyrus, fusiform gyrus; t_max_ = 5.65). The only other task to show significant sdTheta in these regions, to a lesser extent, was the LAT (t_max_ = 2.99).

Overall, the majority of sdTheta occurred in medial and superior frontal cortices, with a right lateralization. LAT and STM were the most widespread in the source space (39% and 35% of significant voxels, respectively), the Game, Music, and PVT intermediate (28%, 27%, 25%), and Speech the least (9%). While most sdTheta sources were frontal, there were substantial differences between tasks. The high theta from the supplementary motor area in the Music task and in the inferior temporal cortex in the Game suggests a preference of sdTheta for cortical areas not critical for the ongoing behavioral task.

### Theta power spectrums during sleep deprivation have multiple peaks

Figure 4 illustrates how the average theta power at baseline more resembles fmTheta in Figure 2A I, especially for the Game, than it does sdTheta within tasks. This suggests that sdTheta occurs in addition to task-related fmTheta found at BL. In order to determine whether sdTheta could be further distinguished from this baseline fmTheta, we inspected the spectrograms of the different tasks for all participants. In particular, we were interested in whether tasks with high frontal BL theta showed an additional distinct peak in the theta range following sleep deprivation. This would support the hypothesis of theta during sleep deprivation as a separate oscillation from task-related, baseline fmTheta.

While statistics at each frequency confirmed that the effect of sleep deprivation was specific to the theta range (Suppl. Figure 5), sdTheta often did not occupy a single consistent peak within or across individuals (Figure 7). Instead, individuals’ peaks were spread over the entire theta range, often with multiple smaller peaks within the same participant. Furthermore, the peak frequency for a given participant was not consistent across tasks (Suppl. Figure 6C-D).

**Figure 7:**
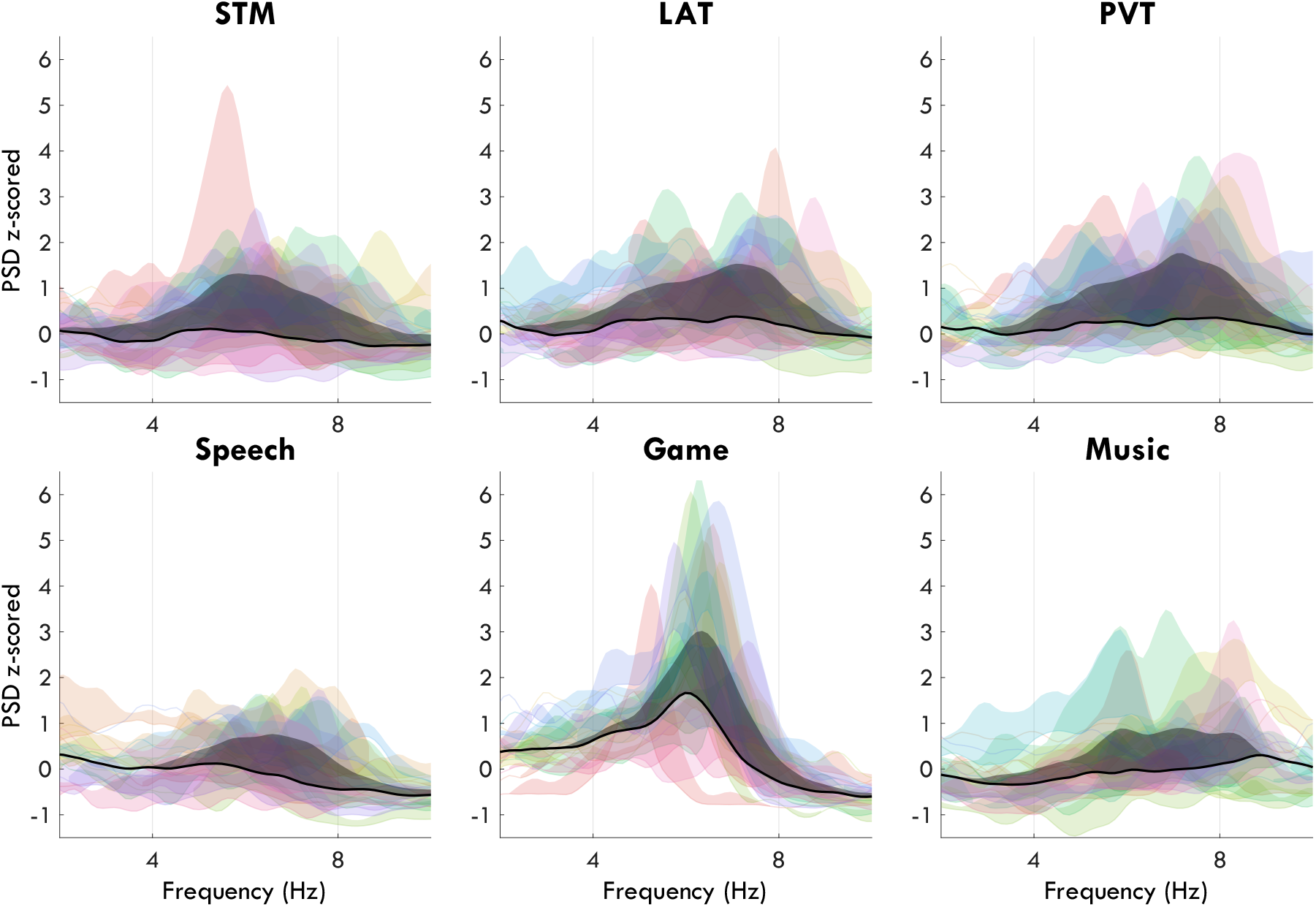
Overlapping z-scored spectrums from the front ROI of each task for every participant. The base curve of each colored patch represents the BL spectrum, the upper curve represents the SD spectrum, and the filled-in area reflects the increase in power. Each participant is a consistent color across tasks. The average power change is the final patch in black. Acronyms: ROI (region of interest), BL (baseline), SD (sleep deprivation), STM (short term memory task), LAT (Lateralized attention task), PVT (psychomotor vigilance task).

The exception was the Game, which showed the overall highest amplitude frontal theta as well as the most clearly defined peak both during BL and SD, with prominence values^2^ (MEAN ± STD) of 1.72 ± 1.14 and 2.94 ± 1.88 respectively. By contrast, the STM task had a prominence of 0.33 ± 0.27 at BL, and 0.71 ± 0.94 at SD (Suppl. Figure 6A-B). In general, the STM had low BL frontal theta, along with Speech and Music (Suppl. Figure 4D).

Due to the clear presence of fmTheta at BL in the Game, we considered this task to be the most likely to show both an fmTheta peak and an sdTheta peak during SD. The BL peak frequency was significantly different from the SD peak frequency, increasing from 5.7 ± 1.0 Hz to 6.4 ± 0.5 Hz (t = 2.62, p = .018, g = 0.84). For reference, the STM peak was 6.0 ± 1.4 Hz at BL, and 6.4 ± 0.7 Hz at SD, but the increase was not statistically significant (t = 0.87, p = .397, g = 0.30). However, as can be seen in the individual Game spectrums in Figure 7, only a single peak is present for most participants, with the baseline theta peak merely shifted in frequency and increased in amplitude during SD. Multiple peaks were instead found in all other tasks during SD, which may indicate a multitude of different theta oscillations not found in the Game.

Visual inspection of the EEG data provided further insight into task-related theta differences. At BL, fmTheta bursts as described by Mitchell et al. (2008) were visible primarily in the Game task (Figure 8A) in 11 individuals. These were frontal-midline bursts that lasted 1-5 s with amplitudes around 15-20 μV. No other prominent theta oscillations were detectable by visual inspection in any task (best example, Figure 8C). During SD, fmTheta became even more prominent in the Game EEG (Figure 8B), with higher amplitudes and longer bursts, appearing for 13 participants and increasing in other tasks as well. In addition to fmTheta, widespread bursts often with frontal peaks appeared during sleep deprivation especially in the LAT and STM (Figure 8D). These had a much shorter duration (2-3 oscillations), but with a higher peak amplitude (> 40 μV). As can be seen from the spectrums (Figure 8 II), game theta bursts yielded narrow-band theta, whereas the LAT bursts had more widespread spectrums. These examples support the interpretation of at least 2 types of oscillations in the theta range that increase with sleep deprivation.

**Figure 8:**
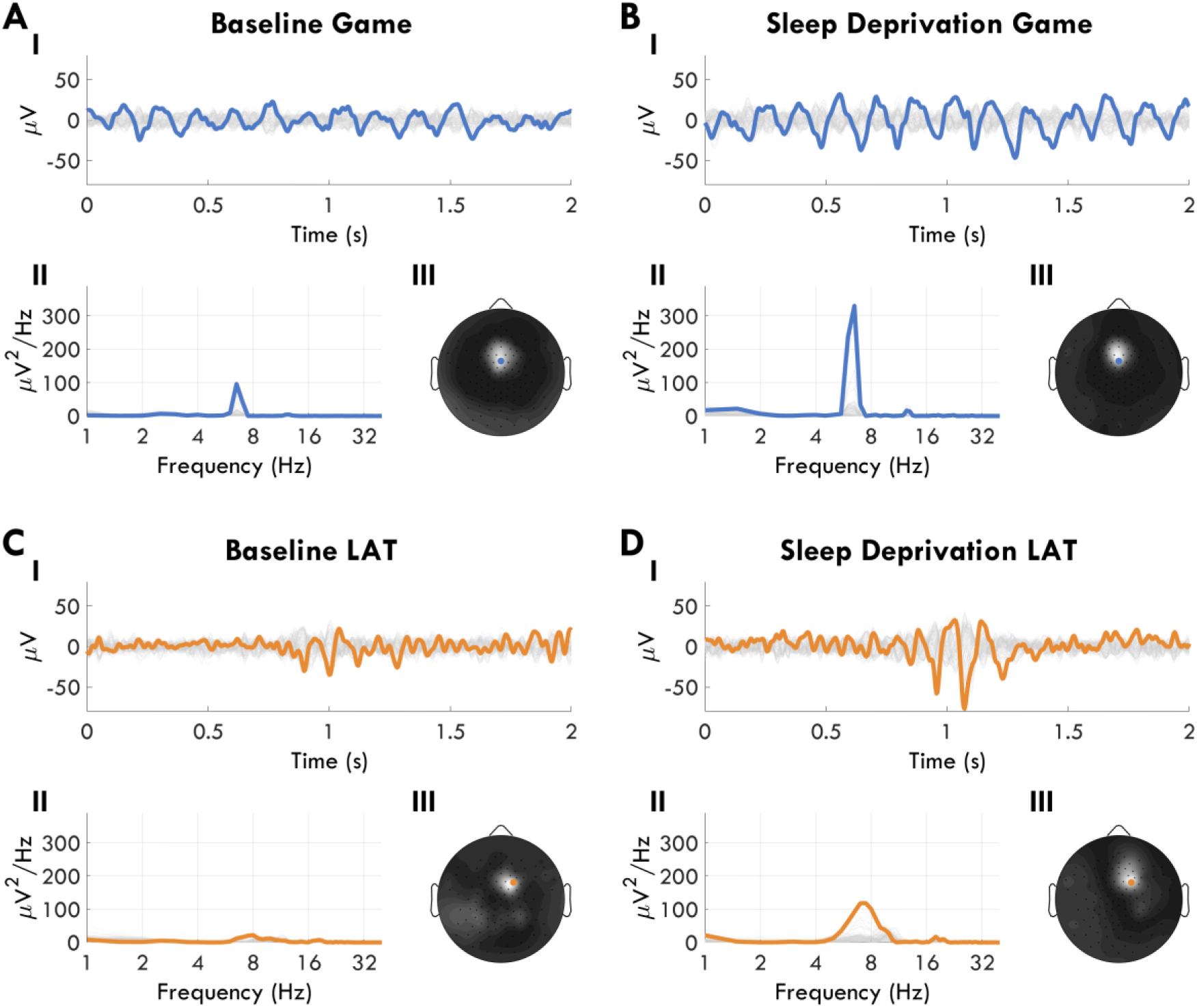
Examples of theta bursts from one participant during BL (**A**, **C**) and SD (**B**, **D**), taken from the Game (**A**, **B**) and the LAT (**C**, **D**). **I**: EEG data in time, amplitude in microvolts. All channels are represented in gray, and the channel expressing the highest theta in color. **II**: Power spectrums of all channels in gray, and peak theta channel in color. X-axis is log-transformed. **III**: Average theta power mapped across all channels from the 2 s shown in I. The scale is normalized for each plot separately to the min-max. Colored dot indicates the same channel highlighted in I and II (ch6 for Game, ch118 for LAT). Acronyms: LAT (Lateralized attention task), BL (baseline), SD (sleep deprivation).

## DISCUSSION

In the literature, there exists two opposing interpretations of theta oscillations: one posits that they reflect a component of cognition, the other that they reflect local sleep. With this study, we investigated whether this apparent paradox could be resolved by the existence of separate oscillations in the theta band, distinguishable by either cortical sources or frequencies. Our results clearly indicate that theta caused by sleep deprivation is not strictly a manifestation of classic fmTheta because: A) their primary sources are in different cortices, namely the right superior frontal gyrus for sdTheta and the left anterior cingulate cortex for fmTheta; and B) sdTheta is present in a much broader subset of cortical areas (Figure 2).

Despite this difference in sources, we did not find distinct theta peaks in the EEG power spectrum (Figure 7) which would have further supported an interpretation of two types of theta oscillations. In Vyazovskiy et al. (2005), sdTheta in rats was manifestly at a lower frequency than the dominant wake hippocampal theta rhythm (5.5 Hz vs 7.5 Hz), with both peaks present during sleep deprivation. This was not replicated in our Game condition, where instead it appears that fmTheta itself increased in amplitude and frequency. For all other tasks, sdTheta occupied a broad range with multiple peaks.

The spectral differences between the narrow-band theta in the Game and broad-band sdTheta in all other tasks could be explained by the different waveforms visually identified (Figure 8): long steady trains of theta in the Game, and high amplitude irregular short bursts in other tasks. These morphological differences make the theta trains comparable to occipital alpha bursts, and the short bursts more comparable to isolated slow waves in sleep. In fact the latter (Figure 8D) would be the most promising candidate for local sleep, compatible with theta events found in rats (Vyazovskiy et al., 2011). The increase in fmTheta in the Game could be a compensation mechanism to counteract the detrimental effect of sleep deprivation, similar to the increase in fmTheta with memory load and cognitive demand. These results could mean that sleep deprivation in humans induces two types of changes in theta: an increase in fmTheta when already present at baseline, and the appearance of local sleep.

An alternative, simpler explanation is that theta may reflect the same mechanism during both cognition and sleep deprivation, regardless of waveform. Simultaneous EEG-fMRI studies previously found that fmTheta originating from the medial prefrontal cortex corresponds to BOLD deactivations in these areas, both during passive rest (Scheeringa et al., 2008) and increasing short term memory load (Scheeringa et al., 2009). Our source localization of sdTheta across the different tasks also suggests that these oscillations may be a marker for cortical areas *not in use*.

First, we found high sdTheta activity in the bilateral (but especially left) supplementary motor area in the Music listening condition. It is compelling that the one task not requiring movement showed such strong theta activity in brain areas involved in complex motor planning (Goldberg, 1985). Notably, the PVT also showed strong activity in bilateral supplementary motor areas, which may seem contradictory. However, the PVT required simply pushing a button after a very obvious stimulus appeared; this made for an almost reflexive response, with little need for deliberative action. By contrast the LAT, which had identical motor requirements but difficult to detect stimuli, despite otherwise widespread high amplitude theta, showed less activity in the supplementary motor areas than the PVT (Figure 6). Supporting this distinction between reflexive and deliberative action, mean reaction times of the LAT were ~20% slower than during the PVT (Suppl. Figure 7B-C), despite identical task requirements (respond within 0.5 s). Vice versa, supplementary motor area activity did not significantly increase in the Speech or Game, two tasks characterized by deliberative action.

Second, high sdTheta was found in the right inferior temporal cortex in the Game, extending all the way to the fusiform gyrus. These areas collectively form the *ventral visual pathway* responsible for object recognition (Ishai et al., 1999). This is in opposition to the *dorsal visual pathway* running from the occipital cortex to dorsal parietal areas such as the supramarginal gyrus and parietal sulcus, where object location is processed (Freud et al., 2016). The Game was almost exclusively a spatial task, requiring participants to map out a target path for a bouncing ball. The only other task to show significant activity in the inferior temporal cortex was the LAT, a spatial attention task. Instead the STM, in essence an object recognition task, showed no significant increase in these areas.

One possible interpretation for theta in cortical areas not in use is that it has a role in *active* inhibition. Such a hypothesis has already been proposed for theta during cognition. Buzsáki in 1996 suggested that theta oscillations in the hippocampus could act as a low-energy solution to selective inhibition (Buzsáki, 1996; Thompson & Best, 1989), such that only neurons synchronized to fire at the correct phase of an ongoing oscillation would successfully transmit action potentials. The role of theta phases in inhibition was supported by phase-targeted closed loop stimulation in mice (Siegle & Wilson, 2014). It may therefore be the case that fmTheta and sdTheta in humans also reflect a low-energy active inhibitory state that conflicting brain networks enter to compensate for cognitive load and sleep deprivation, respectively.

Alternatively, theta could reflect *passive* cortical disengagement. In this scenario, an entire network or brain area ceases to receive inputs, and essentially goes in “standby”. This is comparable to alpha oscillations in visual areas during eyes closed (Kirschfeld, 2005), and may even be related to the bistable default state in which cortical networks enter when physically disconnected from the rest of the brain or during anesthesia (Sanchez-Vives et al., 2017). An interpretation of theta as disengagement, more so than inhibition, would also explain theta activity occasionally found in NREM1 (Santamaria & Chiappa, 1987), at the transition between wake and sleep. In essence, theta as inhibition would be a compensation mechanism for sleep deprivation, whereas theta as disengagement would be a consequence of sleep deprivation, bringing the brain closer to true sleep.

In our results, theta in the supplementary motor areas during Music listening and the inferior temporal cortex in the Game could be interpreted as either active inhibition or passive disengagement. However, the widespread theta increases in the low memory load trials of the STM task relative to the local theta increases in the medium memory load trials (Figure 3C) is more compatible with disengagement during less demanding conditions. Likewise, theta was more widespread in the LAT and STM, subjectively considered more boring tasks (Suppl. Figure 1C), compared to the Speech and Game tasks. However, different experimental methodologies (especially intracortical data) are needed to confirm this link with local neuronal inactivity, and to determine the distinction between active inhibition or passive disengagement.

If fmTheta and sdTheta reflect the same neural process, even cortical disengagement, it becomes more difficult to consider theta as a marker for local “sleep”. The fact that theta is reliably present during rested conditions, especially during complex tasks, speaks against this. The fact that the amplitude of theta power is dependent on local sleep pressure and neural plasticity (Finelli et al., 2000; Hung et al., 2013) is not sufficient to declare these oscillations sleep. In our view, to definitively consider theta events in wake as sleep, especially some form of local slow wave sleep, they would have to show a similar restorative function. If theta is not local sleep but still reflects the same process during cognition, sleep deprivation, and NREM1, then theta could simply reflect local inactivity independent of vigilance state.

Regardless of what theta oscillations truly are, we have demonstrated that both sleep deprivation and tasks can simultaneously affect theta power, and their effects are not easily disentangled. Crucially, increasing sleep pressure resulted in the progressive disappearance of memory load effects of fmTheta, likely due to the higher increase in sdTheta in easier trials. Vice versa, the condition with the overall highest frontal theta was the Game during sleep deprivation, which is also when participants felt, out of all tasks, most awake (Figure 1). As such, theta cannot be used in absolute terms as an objective marker for drowsiness.

Moving forward, research in the EEG of cognition and sleep pressure should not continue to operate blind to each other. Experiments investigating cognitive changes in theta should aim for constant, low levels of sleep pressure. An experiment conducted in the morning may yield different results from an experiment conducted in the evening, or patients who have comorbid sleep disturbances may not show the same theta effects as healthy controls, even with intact cognition. Vice versa, experiments trying to quantify sleep deprivation need to avoid participants arbitrarily increasing their theta by engaging in meditation-like activity, or other forms of focused attention. In general for such studies, it may be more appropriate to record sdTheta during a task like the PVT, which is often used in sleep deprivation studies and simultaneously provides behavioral data. Given the already high variability in theta power across individuals (Suppl. Figure 2), it would be best to reduce variability in mental state. Similarly, we would recommend the use of an “addictive” game to study fmTheta when possible, as this is both enjoyable for the participants and produces remarkably reliable and robust oscillations, more so than the standard short-term memory task.

While our study offers unique insight into theta under different conditions within the same participants, it also suffers limitations. First and foremost, because the participants are from such a narrow age group, these results cannot be generalized to other populations. Theta both during wake and sleep is known to undergo substantial changes across ages, with some theta oscillations disappearing in adolescence and others re-appearing in old age (Ebersole & Pedley, 2003). Additionally, while we mainly interpret these sdTheta results as originating from time spent awake based on previous literature (Cajochen et al., 2002; Finelli et al., 2000), it is likely that they are also affected by circadian fluctuations. Figure 4 illustrates how the frontal spot in particular is only prominent in the SD condition and not SR; this may be due to either the “extreme” sleep pressure or additional circadian components. Furthermore, given the lack of structural MRIs and digitization of electrode positions, the source localization results need to be taken with some caution. It is possible that deeper brain sources were behind these diffuse theta increases, or some other bias we did not account for. Simultaneous EEG-fMRI experiments could possibly resolve these issues. Furthermore, by looking at theta power averaged over several minutes, we do not know if the oscillations appeared synchronously in this network of areas, or in isolation. It is therefore imperative to verify and expand these results with other recording methods, analyses, and more targeted tasks.

In conclusion, we do not provide a definitive resolution to the theta paradox but suggest three possible explanations for our results: 1) fmTheta and sdTheta are separate oscillations, but both can occur during sleep deprivation, one as a compensation mechanism, the other as local sleep; 2) sdTheta is merely a more widespread form of fmTheta, and both reflect active cortical inhibition of task-irrelevant networks; 3) both reflect passive cortical disengagement. EEG offers a limited view of the brain but makes up for it by being easy to use and readily applied in clinical populations and children of any age. As such, it becomes essential to understand as much as possible of the EEG signals we can actually measure. Since its inception, clinicians have been able to visually recognize peculiar oscillations, but such an approach does not lend itself to quantification. Vice versa, analyses such as ours in the spectral domain allow rapid quantification, but at the cost of oversimplification. The results of this study are a reminder that there is still much more to be learned from even basic oscillations like theta.

## MATERIALS & METHODS

### Participants

Participants were recruited from Swiss universities through online advertisements and word of mouth. The advertisements directed students to an online screening questionnaire which if passed provided the participant with the contact information of the experimenters. Detailed screening criteria are provided in the supplementary material. Briefly, participants had to be between 18-25 years old, completely healthy, neurotypical, good sleepers, and at least somewhat vulnerable to sleep deprivation. Due to scheduling restraints caused by the COVID-19 pandemic, some leniency was allowed for edge cases (e.g. one participant was 26 at the time of recording).

75 applicants conducted the screening questionnaire. One was recruited for technical pilots (data not included), and an additional 31 passed but did not initiate contact or were unable to meet the scheduling requirements. 19 participants were recruited, however one dropped out midway and so was not included in further analyses. Of the 18 participants who completed the experiment, 9 were female and 3 were left-handed. Mean age (± standard deviation) was 23 ± 1 years old. All participants self-reported above-average English fluency (68% ± 13% on a scale from *terrible* to *native speaker*), with 1 participant a native English speaker. All had corrected-to-normal vision and self-reported no hearing impairments.

Data collection and interaction with participants was conducted according to Swiss law (Ordinance on Human Research with the Exception of Clinical Trials) and the principles of the Declaration of Helsinki, with Zurich cantonal ethics approval BASEC-Nr. 2019-01193. All participants signed informed consent prior to participation and were made aware that they could terminate the experiment at any time.

### Experiment design

Participants came to the laboratory twice, first for the baseline then the sleep deprivation recordings, separated by at least 4 days. During the week prior to each session, participants were asked to maintain a regular sleep wake cycle, going to bed and waking up within 1h of a pre-determined sleep and wakeup time based on their personal preference. These individualized sleep and wake times were then used during the experiment. During the control week, participants wore a wrist accelerometer (GENEActiv, Activinsights Ltd.) and filled out regular sleep reports to ensure compliance. Participants were further asked to abstain from alcohol in the 3 days prior to the measurement, and limit caffeine consumption to no more than the equivalent of 2 cups of coffee, and never after 16:00. They were asked to avoid time-zone travel and any activities they knew could affect their sleep (e.g. parties, skiing, sauna).

Data was collected in Zurich, Switzerland, between February and December 2020, overlapping with the COVID-19 pandemic and consequent lockdowns. Due to scheduling restraints, 4 participants conducted the baseline after the sleep deprivation recording, so the experimental session orders were not balanced.

#### Baseline

After the EEG net was set up (for details, see *Supplementary material: Methods: Recording equipment*), participants went to bed at the agreed-upon time (21:55-00:47), and were left to sleep for as long as they wished (6.2-10.3 h). In the morning, participants first filled out a sleep quality questionnaire while the researcher refreshed the EEG gel and restored signal quality. Then, participants were provided breakfast and given time to wake up. Finally, participants performed the baseline (BL) task block (8:10-11:17), 1.8 ± 0.6 h from wake onset. A brief resting wake recording was conducted in the evening and in the morning, however the data was not included in this manuscript. The complete schedule is depicted in Figure 9.

**Figure 9:**
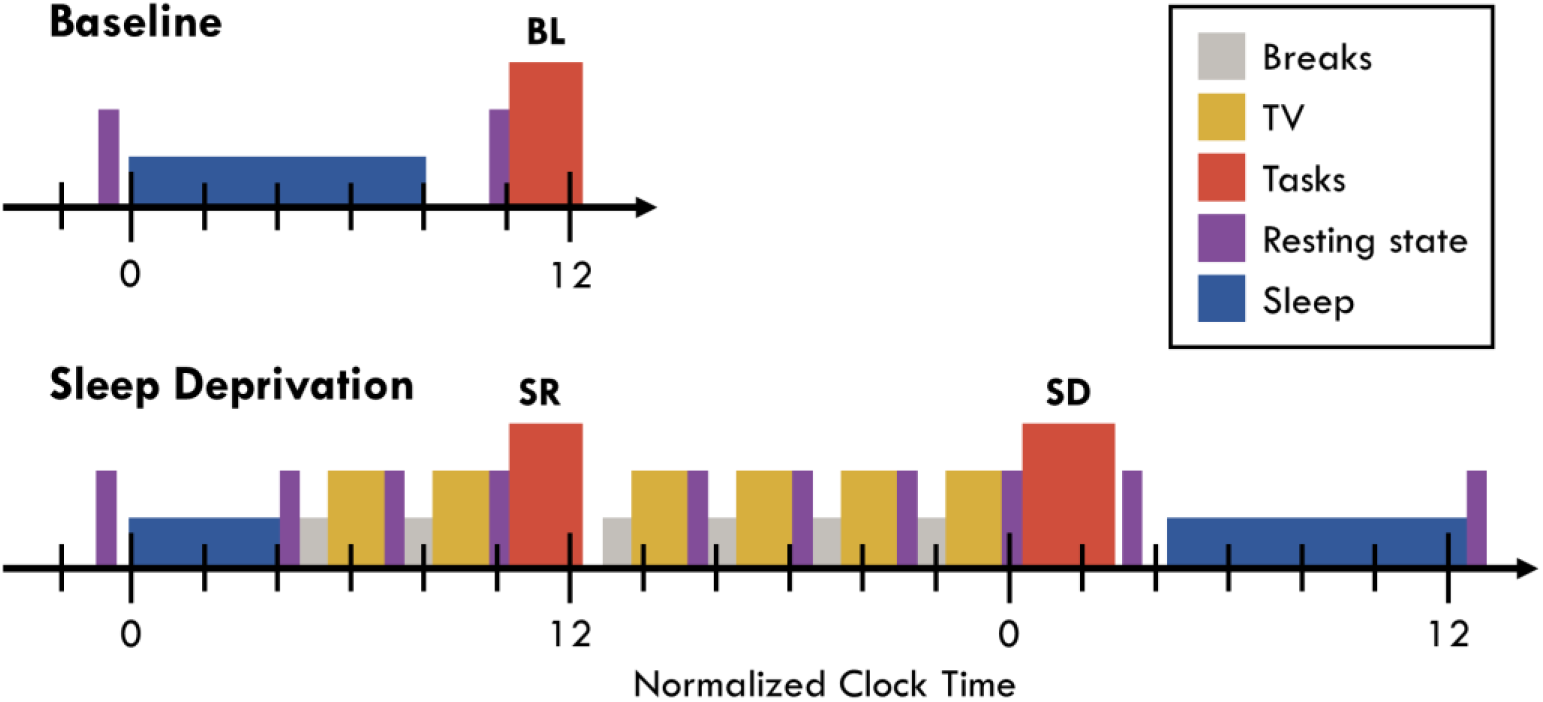
Experiment timeline example for a sleep-wake schedule of 0:00-8:00. Unlike during the control week, participants were free to wake up when they wished at baseline and during the recovery night. This was done to allow for recovery sleep in case of poor sleep quality and ensure participants were fully rested the morning after. Empty spaces were transition periods allowing for delays in the schedule. The SD block was longer than other task blocks because of additional repetitions of the LAT following the main block (data not included). Acronyms: BL (baseline), SR (sleep restriction), SD (sleep deprivation), LAT (lateralized attention task).

#### Sleep deprivation

Participants followed the same evening routine as in the baseline, going to bed at the same time. They were woken up 4 hours later and conducted the same morning routine. Then throughout the day, participants repeated 6 cycles consisting of a break, 2 TV episodes from a series of their choice, and a brief rest recording. During the breaks, participants were provided a small home-cooked meal (selecting items from a menu beforehand), thus eating the same plate during every break. They repeated 2 of these cycles in the early morning, then conducted the morning sleep restriction (SR) task block after 6.4 ± 0.2 h from wake onset (within 7.7 ± 39.5 min of the BL block). Participants went through 4 more cycles before conducting the sleep deprivation (SD) task block, after 20.0 ± 0.1 h from wake onset and within 2.6 ± 10.5 min of the prior night’s bedtime. After 23.6 ± 0.5 h of wake, participants went to bed and slept for as long as they wished. During all wake recordings, participants were monitored by an experimenter to ensure they did not fall asleep. From the evening before the first night to the day after the recovery night, participants remained in the sleep laboratory and did not have access to clocks or external time cues. Two participants reported nausea with increasing sleep deprivation and were therefore provided a break outside just prior to the SD block (in complete nocturnal darkness).

### Tasks

Each task block lasted approximately 2 hours. The order of tasks was randomized and counterbalanced across participants. For each participant, tasks were conducted in the same order for all three blocks. Each task began and ended with a 1 min rest period allowing participants to adjust and get comfortable. After each task, participants answered a task battery questionnaire asking how they experienced the task (Suppl. Figure 1).

#### Short-Term Memory Task (STM)

Participants performed a ~25 min delayed match-to-sample / short term memory task, adapted from Habeck et al. (2004) and Maurer et al. (2015). The task consisted of 120 trials divided in 4 blocks, with 3 levels of memory load randomized across trials for a total of 40 trials per load. Stimuli are depicted in Figure 10A. Each trial was separated by a 1-2 s pause with a black screen, The ENCODING epoch began when a red fixation square appeared in the center of the screen for 1 s. Then 1, 3, or 6 symbols (selected from a pool of 30 “letters” of the Aurebesh fictional alphabet) were displayed around the fixation point in 8 possible locations for 2 s. Participants were instructed to maintain fixation on the red square while memorizing these symbols. This was followed by a 4 s RETENTION window in which only the fixation point was displayed, and participants had to hold in memory the symbols from the encoding epoch. For all analyses, this retention period was divided into two 2 s epochs, and only data from the first epoch was used in this paper. The trial ended with the PROBE epoch, in which a probe symbol replaced the central fixation point and participants had to indicate with left or right arrow keys whether the probe symbol was contained in the encoding set or not within a 3 s window. The probe was from the encoding set in 50% of trials. No feedback on performance was provided.

**Figure 10:**
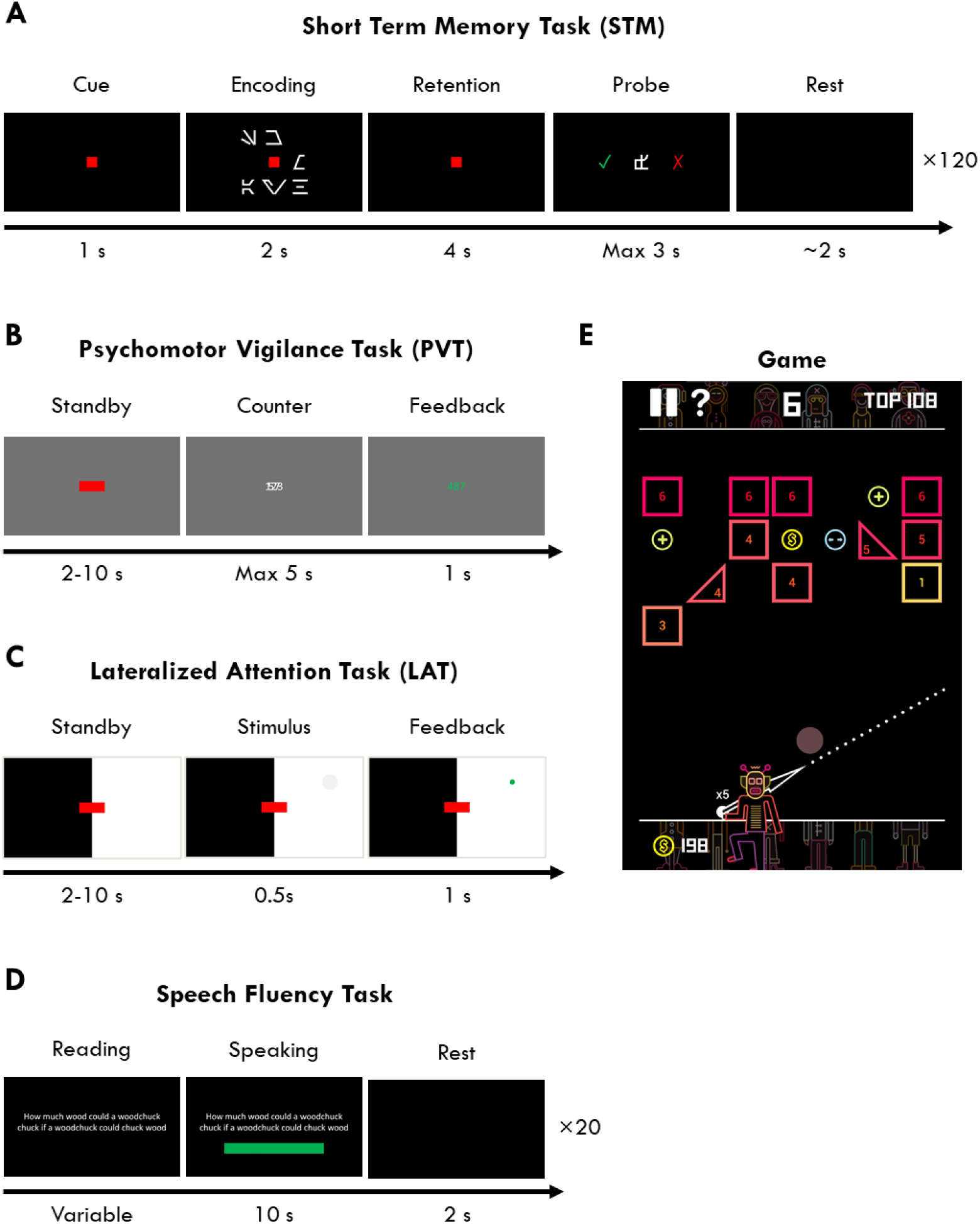
Stimuli used for the tasks.

#### Psychomotor Vigilance Task (PVT)

This is a standard reaction-time task used in sleep deprivation paradigms, based on (Basner & Dinges, 2011). The total task duration was 10 min. Participants were presented with a red fixation rectangle on a gray background (Figure 10B). Every 2-10 s, the rectangle was replaced with a millisecond countdown and participants had to press a button as fast as possible to stop it. The response time would then freeze for 1 s and be colored in yellow if less than 0.1 s (false alarm), green if between 0.1 and 0.5 s (correct response), and red if later than 0.5 s (lapse). If participants did not respond within 5 seconds, an alarm would sound to wake them up.

#### Lateralized Attention Task (LAT)

this was a 12 min visuo-spatial reaction time task, modelled after the PVT. 6 blocks (2 min each) alternated between having the left or right visual hemifield in white, and the other in black (Figure 10C). Participants had to maintain fixation on a red rectangle in the center of the screen, and covertly attend to the white half of the screen. Every 2-10 s a feint grey circle (1cm radius, #F7F7F7) would appear randomly in any location of the illuminated hemifield and shrink away completely within 0.5 s. The circle would freeze and flash green when the button was pressed before it disappeared. If 5 stimuli were missed consecutively, an alarm would sound to wake up the participant. During the delay periods, 50 ms pink noise tones were presented every 1.5-5 s at ~50dB. Participants were instructed to ignore these tones.

#### Speech Fluency Task

Participants performed a tongue-twister reading task in English for 5-10 min. This consisted of 20 trials, one for each sentence. Each trial began with the sentence written on the screen (Figure 10D). Participants were instructed to read it in their head once or twice to get familiar with it, but not practice speaking. When they were ready, they could press a button, and a green bar would appear below, steadily shrinking to count down a 10 s reading window. In this time, participants had to read out loud the sentence as many times as possible, as clearly as possible, and as correctly as possible.

#### Game

Participants played the mobile game BBTAN (by 111%, based on the 1986 game *Arkanoid* by Taito) for 10 minutes (Figure 10E). They started each session from level 1. The game involved a robot with a ball at the bottom of the screen, and a row of 1-6 bricks at the top. By tapping and dragging on the screen, participants could orient an arrow from the robot, and the ball would be launched from the robot in the indicated direction. The goal was to bounce the ball against the walls and hit as many bricks as possible, such that every time the ball hit a brick, the brick lost a point, and when the brick had no more points, it disappeared. At each round, after the ball was launched, hit the bricks, and bounced back to the bottom, the remaining set of bricks descended by 1 row, and a new row of bricks appeared at the top. When the bottom-most row of bricks reached the robot, the player lost the game. There were additional game features to help remove bricks faster. This was a “simple but addictive” game, requiring a minimum amount of spatial strategy to win, without any time pressure.

#### Music

Participants listened to two songs for 2.5 min each: the beginning of the instrumental soundtrack *Light of the Seven* (composed by Ramin Djawadi from *Game of Thrones: Season 6*), and the beginning of the soundtrack *Finale (William Tell Overture*) composed by Hans Zimmer from *The Lone Ranger*.

### EEG analysis

High-density EEG was recorded using HydroCel Geodesic Sensor Nets^™^ with 128 channels, connected to DC BrainAmp Amplifiers and recording software Brainvision Recorder (Vers. 1.23.0003, Brain Products GmbH, Gilching, Germany). Data was recorded with a sampling rate of 1000 Hz with Cz reference.

All data preprocessing, analysis, and statistics was done with custom scripts in MATLAB (R2019b) based on the EEGLAB toolbox v2019.1 (Delorme & Makeig, 2004). All further analyses involving source localization were performed with the FieldTrip toolbox v20210606 (Oostenveld et al., 2011). Detailed analysis pipelines are provided in the supplementary material, and code is available on GitHub (https://github.com/snipeso/Theta-SD-vs-WM).

#### Preprocessing

EEG data was filtered between 0.5-40 Hz and downsampled to 250 Hz. Visual detection of major artifacts and bad channels was conducted by author SS, blind to participant, task, and session. ICA was then used to remove physiological artifacts, mainly eye movements, heartbeat, and muscle activity (Dimigen, 2020). Bad channels were interpolated, and only the first 4 minutes of clean data were used. The full pipeline is depicted in Suppl. Figure 8.

#### Channel space power calculation

120 channels were used. Power source density (PSD) was calculated using MATLAB’s pwelch function, with 8 s windows, Hanning tapered, and 75% overlap. Data for each frequency was z-scored. For theta topographies (e.g. Figure 4), z-scored PSD values between 4-8 Hz were averaged. For power spectrums (e.g. Figure 7), z-scored PSD values were averaged into 3 pre-selected regions of interest (ROIs). Exact channels are available in the Supplementary Material. For mean theta values (e.g. Figure 3B), these ROI spectrum averages were further averaged between 4-8 Hz. The channel space analysis pipeline is depicted in Suppl. Figure 11.

#### Source localization

Beamformer source localization was done with the dynamic imaging of coherent sources (DICS) algorithm from FieldTrip. A finite-element head model was implemented with the SimBio toolbox (Vorwerk et al., 2018) based on the segmentation of a standard MRI template brain. A 3D grid with 10 mm resolution (3294 voxels) was used as a source model. After being projected into the source space, power was z-scored for each frequency. For visualization, t-tests were conducted for all gray-matter voxels, cluster corrected for multiple comparisons, and significant clusters projected onto the inflated brain. To determine the main anatomical sources, z-scored data was parcellated based on the Automated Anatomical Labelling (AAL) atlas (Tzourio-Mazoyer et al., 2002). The median value of all voxels within each area was then averaged across frequencies. For both pipelines, only cortical areas were included, as there is currently little evidence that activity from deep brain structures reaches the scalp. Further details can be found in *Supplementary material: Methods: Source localization*, with the source space analysis pipeline depicted in Suppl. Figure 12.

#### Trial analysis

Power calculations in both channel and source space for the 2 s retention windows of the STM task were done as described above, except using artifact-free non-overlapping trials from the entire 25-minute recording. Trials were first averaged within each session by load level, and then z-scored pooling levels, sessions, and channels for each participant and each frequency. The minimum number of trials for each level for each session was 25.

#### Statistics

All parametric statistics were based on α = 5%. For both the channel space and source localized anatomical areas, paired t-tests were conducted between BL and SD. False discovery rate (FDR) correction was then applied for each comparison (Benjamini & Hochberg, 1995). When t-values are specified in the text, FDR corrected p-values are provided, along with Hedge’s g effect sizes (Hentschke & Stüttgen, 2011). One PVT BL recording is missing, otherwise there were always 18 datasets per task, per session. Further details can be found in *Supplementary material: Methods: Statistics*.

## ACKNOWLEDGEMENTS

This study was conducted as part of the SleepLoop Flagship project of Hochschulmedizin Zürich, with additional funding from the Swiss National Science Foundation (320030_179443) and Hirnstiftung. Professor Hans Peter Landolt provided the sleep laboratory, Professor Christian Baumann and Niklas Schneider the EEG equipment. Simone Accascina helped create the EWOQ platform, task programming, and provided technical support during data collection. Tommaso Fedele provided consultation for the source localization. Noa Rieger aided with data collection and Selina Schuele conducted the sleep scoring. Lastly, a special thank you to all of our participants for taking part in this study.

## Footnotes

1 When tmax is specified, then the highest t-value from neighbouring or group of channels/areas is used

2 Difference in z-scores between the maximum theta amplitude and the closest trough in the spectrum

## Notes

### Competing Interest Statement

The authors have declared no competing interest.

