## Supplementary Material for "The theta paradox: 4-8 Hz EEG oscillations reflect both local sleep and cognitive control"

#### RESULTS

##### Subjective ratings after each task

A brief questionnaire was conducted after each task to assess participants' subjective experiences. All answers were provided on a continuous scale. A two-way rmANOVA was conducted for each question to assess the effects of *session*, *task*, and their interaction (Table 1). All means are shown in Figure 1.

Subjective sleepiness demonstrated by far the largest effect of session. All other questions showed modest effects of session (.019 < $\eta^2$ < .070).

| | Session | Task | Interaction |
|---|---|---|---|
| Subjective sleepiness | $F_{(2, 30)}$ = 35.42, **p < .001**, $\eta^2$ = .355 | $F_{(5, 75)}$ = 14.7, **p < .001**, $\eta^2$ = .073 | $F_{(10, 150)}$ = 0.96, p = .440, $\eta^2$ = .008 |
| Relaxing | $F_{(2, 30)}$ = 5.13, **p = .012**, $\eta^2$ = .019 | $F_{(5, 75)}$ = 41.97, **p < .001**, $\eta^2$ = .530 | $F_{(10, 150)}$ = 1.11, p = .365, $\eta^2$ = .010 |
| Engaging | $F_{(2, 30)}$ = 6.42, **p = .015**, $\eta^2$ = .027 | $F_{(5, 75)}$ = 43.63, **p < .001**, $\eta^2$ = .548 | $F_{(10, 150)}$ = 1.52, p = .191, $\eta^2$ = .010 |
| Focus | $F_{(2, 30)}$ = 5.25, **p = .016**, $\eta^2$ = .039 | $F_{(5, 75)}$ = 13.63, **p < .001**, $\eta^2$ = .235 | $F_{(10, 150)}$ = 0.39, p = .853, $\eta^2$ = .006 |
| Difficulty | $F_{(2, 30)}$ = 7.02, **p = .006**, $\eta^2$ = .038 | $F_{(4, 60)}$ = 31.67, **p < .001**, $\eta^2$ = .426 | $F_{(8, 120)}$ = 0.71, p = .585, $\eta^2$ = .008 |
| Effort | $F_{(2, 28)}$ = 2.17, p = .139, $\eta^2$ = .022 | $F_{(4, 56)}$ = 4.84, **p = .015**, $\eta^2$ = .114 | $F_{(8, 120)}$ = 0.37, p = .838, $\eta^2$ = .006 |
| Performance | $F_{(2, 30)}$ = 2.02, p = .156, $\eta^2$ = .025 | $F_{(4, 60)}$ = 8.96, **p < .001**, $\eta^2$ = .134 | $F_{(8, 120)}$ = 1.04, p = .401, $\eta^2$ = .023 |
| Motivation | $F_{(2, 20)}$ = 6.93, **p = .021**, $\eta^2$ = .070 | $F_{(5, 50)}$ = 19.61, **p < .001**, $\eta^2$ = .381 | $F_{(10, 100)}$ = 3.33, **p = .020**, $\eta^2$ = .04 |

***Table 1:*** *2-way repeated measures ANOVA with factors* session, task *and their interaction for every subjective rating. Significant p-values are in bold (not correcting for multiple comparisons). N.B. not all questions were answered by all participants, and therefore there are differences in the degrees of freedom.*

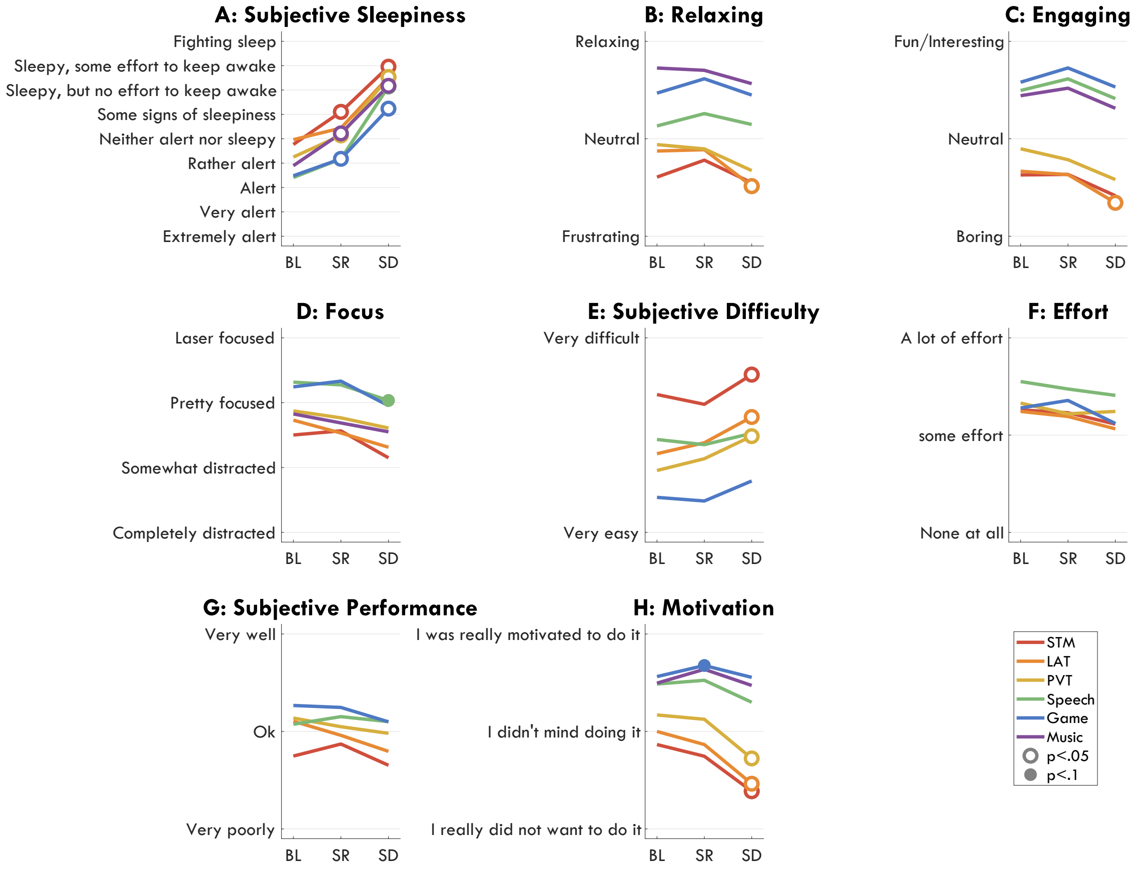


***Figure 1:*** *Questionnaire answers following each task during each session. All answers were given on a ~10 cm slider with labels at specific intervals (indicated on the y-axis). White circles indicate a significant change from BL, filled circles a trend, FDR corrected. Each question was asked in alphabetical order.* ***A****: "Please indicate your sleepiness right now." Adapted from the Karolinska Sleepiness Scale (KSS).* ***B-C:*** *"How did you experience this task?"* ***D:*** *"How focused on the task were you?"* ***E****: "How hard was it to perform this task?"* ***F****: "How much effort did you put into performing this task? (Think about how much you tried to do well, and how much more you could have done)."* ***G:*** *"How well do you think you did the task?"* ***H:*** *"How motivated were you during the task?". Acronyms: STM (short term memory task), LAT (lateralized attention task), PVT (psychomotor vigilance task), BL (baseline), SR (sleep restriction), SD (sleep deprivation), FDR (false discovery rate).*

#### Extreme individual differences in theta power

Raw power amplitudes were extremely variable across participants, especially within the 4-8 Hz range. Z-scoring for each frequency (Main Figure 7) allowed for all participants' spectrums to occur within the same range, and highlighted more accurately the systematic changes across sessions, tasks, and channels. Notably, while the untransformed spectrums identified only 3-4 participants with increased theta with SD (Figure 2), the same z-scored values (Main Figure 7) revealed a theta increase in essentially all participants.

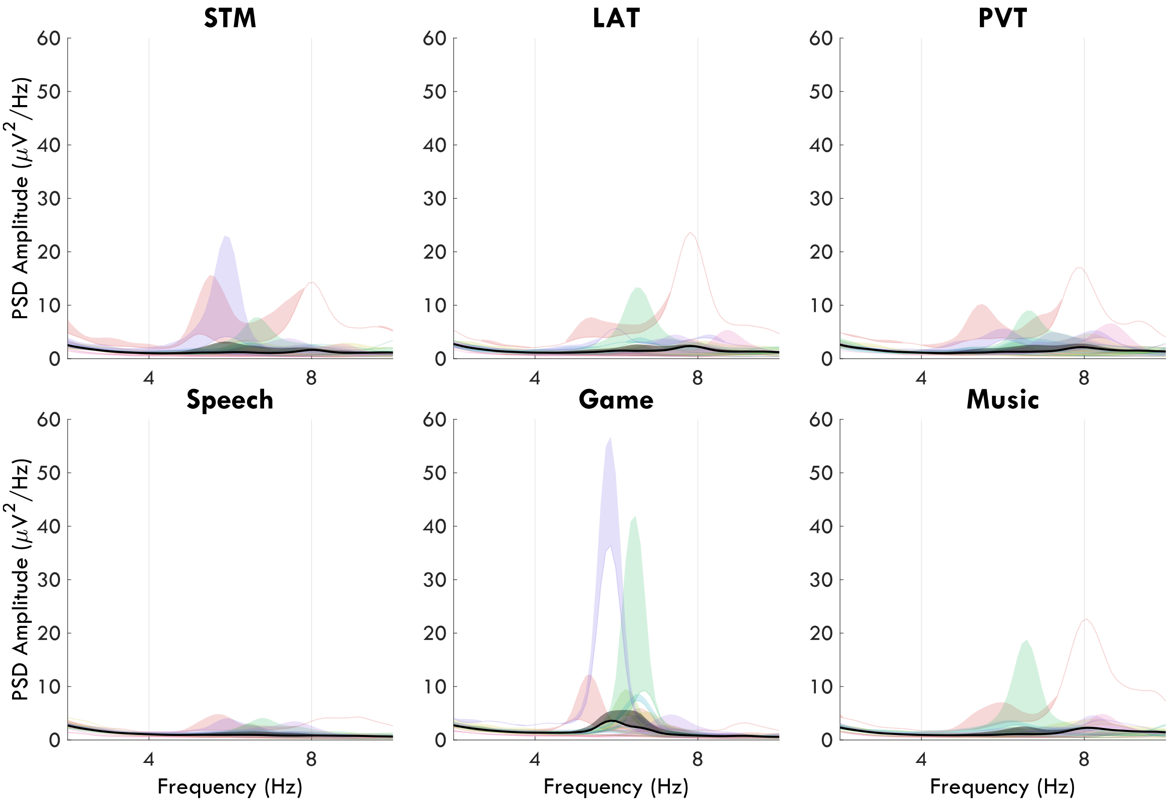


***Figure 2:*** *Overlapping EEG power spectrums, untransformed, from the first 4 min of the front ROI of each task for every participant. Each participant is a consistent color across tasks. The base curve of each colored patch represents the BL spectrum, the upper curve represents the SD spectrum, and the filled-in area reflects the increase in power. The average power change is the final patch in black. Acronyms: EEG (electroencephalogram), ROI (region of interest), PSD (power source density), BL (baseline), SD (sleep deprivation), STM (short term memory), LAT (lateralized attention task), PVT (psychomotor vigilance task).*

#### ROI spectrums for the STM retention period

Figure 3 provides the z-scored average spectrums for the three ROIs (front, center, back) for the first 2 s retention period of the short-term memory (STM) task, highlighting the differences across memory load for each session. Notably, L3 theta has a double peak at BL in the 4-8 Hz range, whereas all levels have comparable frontal spectrums during SD. Given the double peak and the low frequency resolution for 2 s windows (0.5 Hz), these spectra were not used in the manuscript to determine whether fmTheta had a different peak frequency than sdTheta.

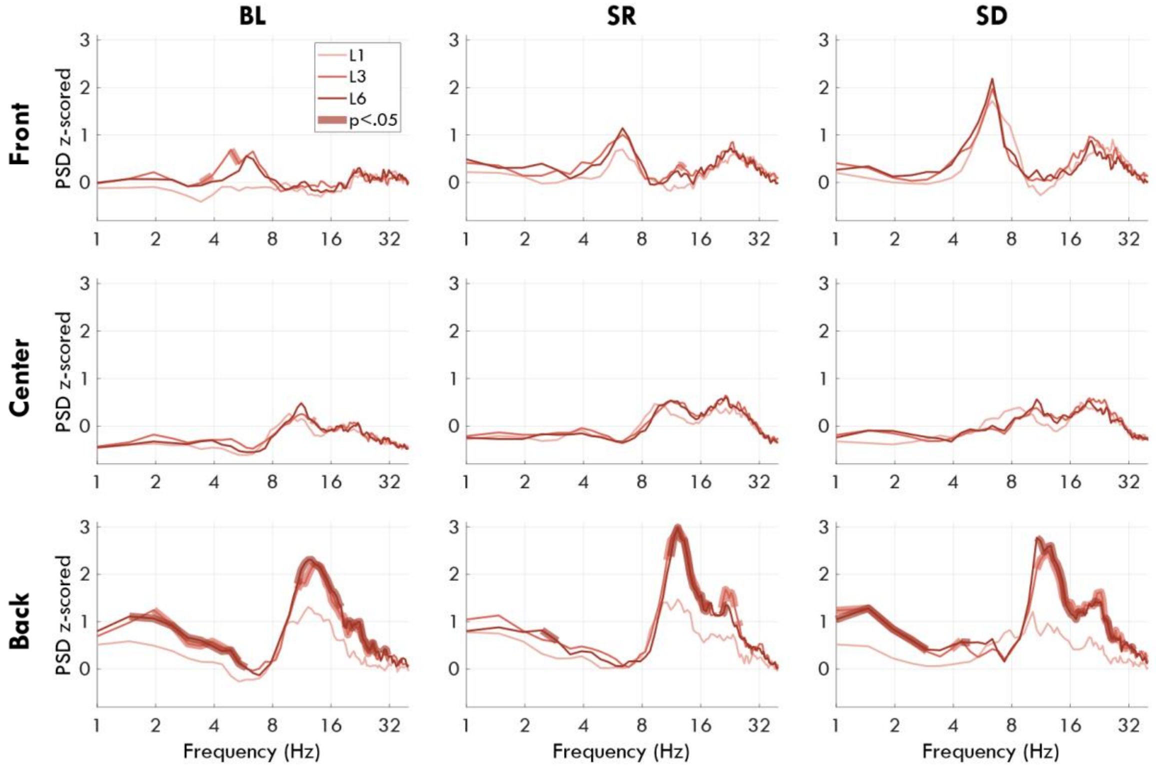


***Figure 3:*** *Spectrum of the first half of the retention period (2 s) of the STM task. Each row represents a different ROI, each column a different session. The thin lines represent the spectrums for the three memory loads (1, 3 and 6 items). Thick lines represent frequencies that were significantly different from L1, FDR corrected. The x-axis is log-transformed. Acronyms: PSD (power spectral density), STM (short term memory task), BL (baseline), SR (sleep restriction), SR (sleep restriction), SD (sleep deprivation), FDR (false discovery rate), ROI (region of interest).*

#### Average theta in ROIs

| | Session | Task | Interaction |
|---|---|---|---|
| Front | $F_{(2, 32)} = 28.02$, $\mathbf{p < .001}$, $\eta^2 = .224$ | $F_{(5, 80)} = 22.51$, $\mathbf{p < .001}$, $\eta^2 = .249$ | $F_{(2, 160)} = 1.88$, $p = .090$, $\eta^2 = .010$ |
| Center | $F_{(2, 32)} = 13.09$, $\mathbf{p < .001}$, $\eta^2 = .105$ | $F_{(5, 80)} = 14.05$, $\mathbf{p < .001}$, $\eta^2 = .239$ | $F_{(2, 160)} = 2.53$, $\mathbf{p = .035}$, $\eta^2 = .021$ |
| Back | $F_{(2, 32)} = 2.41$, $p = .111$, $\eta^2 = .021$ | $F_{(5, 80)} = 21.67$, $\mathbf{p < .001}$, $\eta^2 = .305$ | $F_{(2, 160)} = 0.79$, $p = .549$, $\eta^2 = .007$ |

***Table 2:*** *2-way rmANOVAs for z-scored theta power changes for 3 regions of interest (ROI) across all sessions (BL, SR, SD) and tasks (STM, LAT, PVT, Music, Game, Speech).*

Overall, the effects of task and sleep deprivation were similar. In the front ROI (Figure 4A), sleep deprivation substantially increased theta to a similar extent in all tasks, however the effect was highest in the PVT and lowest in the Music and Speech tasks (Figure 4E). The Game had the highest overall frontal theta, the LAT and PVT had intermediate levels, and the STM, Music, and Speech tasks had the lowest (Figure 4D). Notably, theta did not continue to increase from SR to SD in the center ROI for the Game, Speech, and Music tasks (Figure 4B), driving the significant interaction in the 2-way rmANOVA.

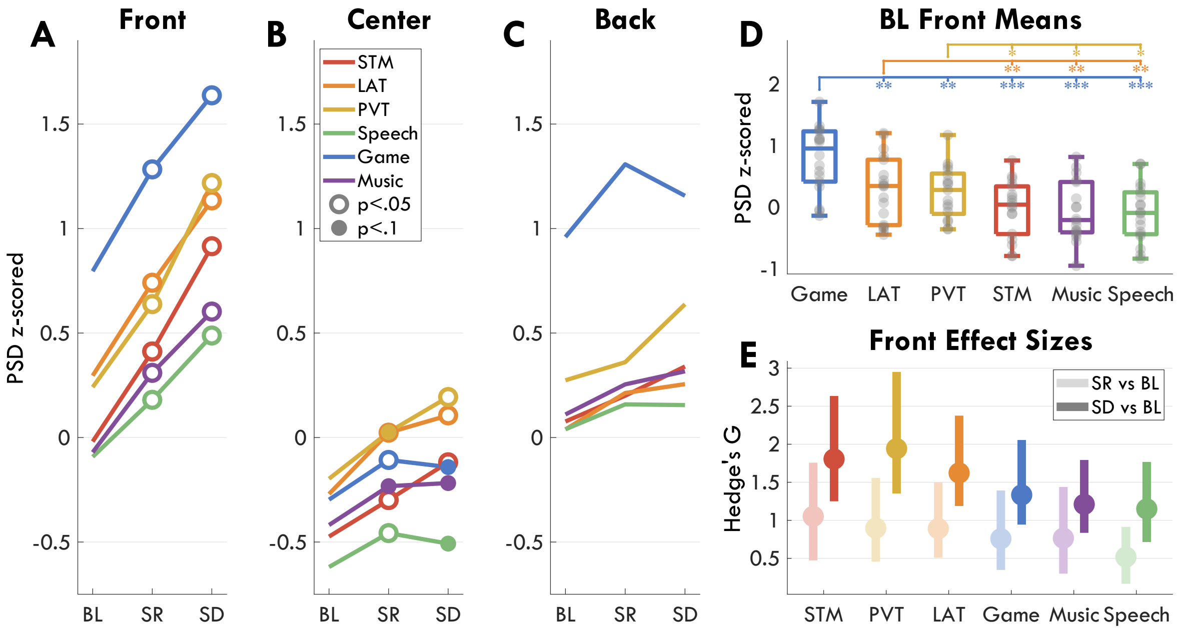


***Figure 4****: Mean z-scored theta power by task across sessions for 3 ROIs: Front (**A**), Center (**B**) and Back (**C**). Open circles indicate within each task a significant change in theta relative to BL, filled circles indicate a trend, based on paired t-tests, FDR corrected for multiple comparisons within each plot.* ***D****: Mean theta power for all tasks at baseline in the front ROI. Gray circles represent each participant, the boxplot indicates median and interquartile range. Stars indicate significant differences between tasks (the color indicates one task, the location of the stars the other) such that: * p-value < .05, ** p-value < .01, *** p-value < .001.* ***E****: Hedge's g effect sizes of the changes in theta in the front ROI from BL to SR (light colors) and SD (dark colors). The disk indicates the measured g value, the bars indicate 95% confidence intervals. Tasks are ordered by g values. Acronyms: ROI (region of interest), PSD (power spectral density), STM (short term memory task), LAT (lateralized attention task), PVT (psychomotor vigilance task), BL (baseline), SR (sleep restriction), SD (sleep deprivation), FDR (false discovery rate).*

#### Power increase with sleep deprivation is specific to the theta and beta ranges

To determine the specificity of the changes in theta power across sessions to the 4-8 Hz range, we compared the change in each frequency (1-35 Hz) for each ROI during each task across sessions using paired t-tests. Results are shown in Figure 5. Overall, the largest changes with session were in the theta and beta (15-25 Hz) ranges, primarily in the front ROI, in line with previous literature (Cajochen et al., 2002; Finelli et al., 2000; Strijkstra et al., 2003). Given the lack of change in the delta (1-4 Hz) and alpha (8-12 Hz) ranges, the effects in the main manuscript cannot be attributed to either slow waves or alpha bursts, nor a broadband increase in power.

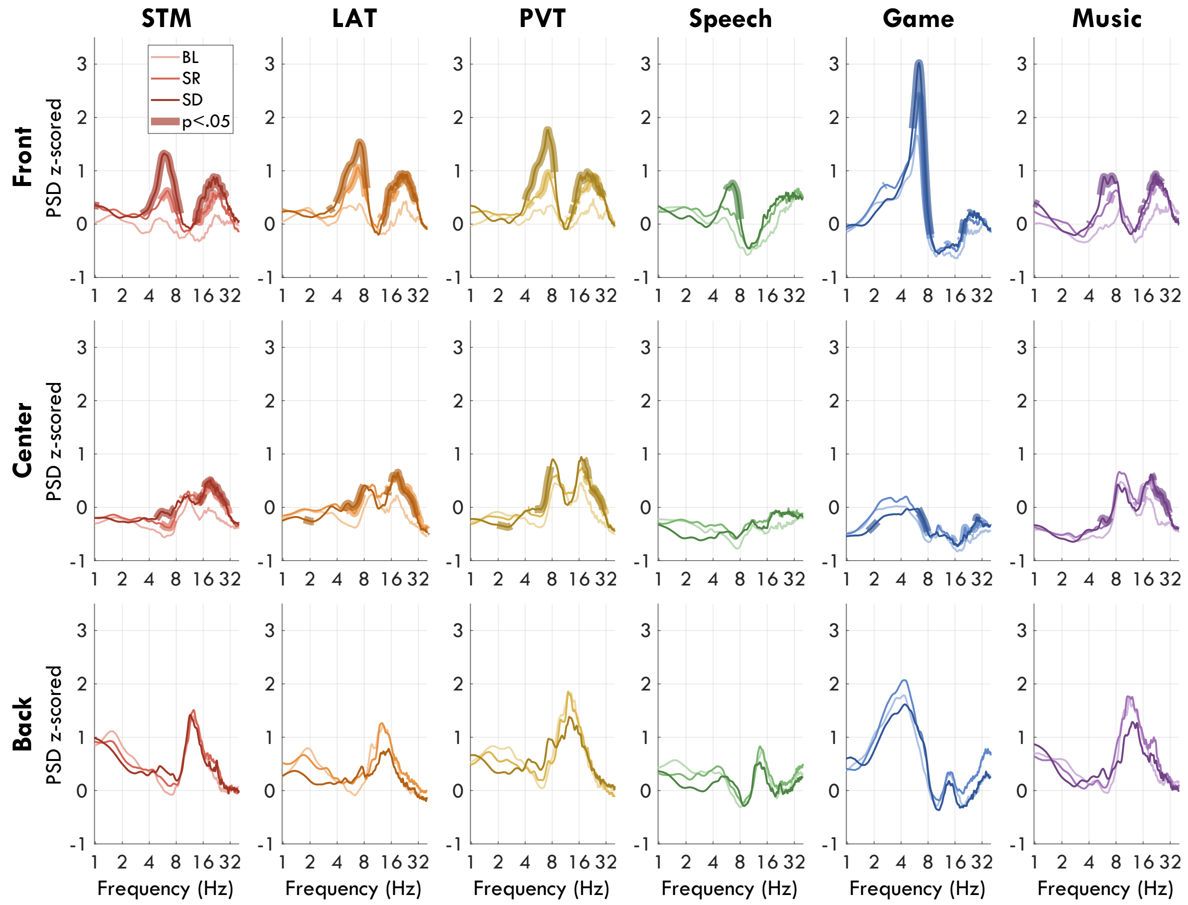


*__Figure 5:__ Power spectrums for each ROI during each task. Thin lines indicate the spectrum at each session. Thick lines indicate statistically significant changes (paired t-tests, p-value < .05, FDR corrected) for a given frequency relative to BL. The frequency axis is log-transformed. The y-axis represents z-scores. Acronyms: ROI (regions of interest), PSD (power spectral density), STM (short term memory task), LAT (lateralized attention task), PVT (psychomotor vigilance task), BL (baseline), SR (sleep restriction), SD (sleep deprivation), FDR (false discovery rate).*

#### Theta peak properties

In order to claim that fmTheta and sdTheta differ by spectrum, 3 conditions should be met: 1) sdTheta and fmTheta have sufficiently (e.g. ≥ 0.5 Hz) different peak frequencies; 2) these peak frequencies are actually from a defined peak; 3) both peaks are present during a cognitive task at SD. Peak frequency was measured as the highest peak in the 3-9 Hz range. Prominence was quantified as the change in amplitude from this peak to the nearest trough in the spectrum; if there was only one peak, then it became the difference between minimum and maximum power. Figure 6A indicates how only the Game had a prominent theta peak at BL and maintained the relatively higher peak during sleep deprivation. This makes the Game task the only candidate to properly identify both fmTheta and sdTheta. Complicating further the analysis of peaks, Figure 6C-D illustrates how even within individuals, theta peak frequencies were not constant across tasks.

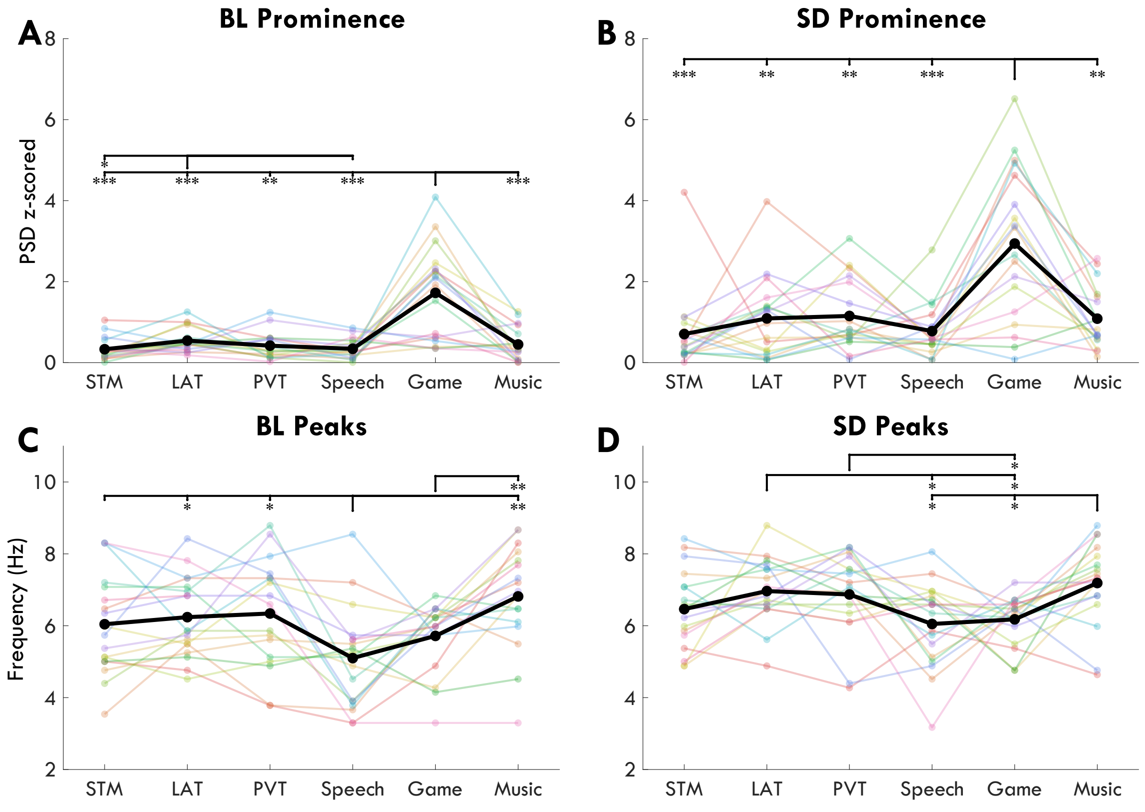


***Figure 6:*** *Prominence and peak frequency of z-scored power spectrums between 3 and 9 Hz for each task. Each color represents a different participant, the black line indicates the average. Asterisks indicate significant differences from paired t-tests between tasks, FDR corrected, such that: * p-value < .05, ** p-value < .01, *** p-value < .001.* ***A-B****: Prominence refers to the amplitude difference between the highest peak and the closest trough to that peak within a 3-9 Hz range.* ***C-D****: "Peaks" refers to the frequency of the highest amplitude peak in the 3-9 Hz range.*

#### Behavioral results

The impact of sleep deprivation on behavioral performance has already been addressed extensively by others (Lim & Dinges, 2010; Lo et al., 2012), finding vigilance to be the main cognitive function affected by acute sleep deprivation. Here, we provide the performance measures of the STM, LAT, PVT, and Speech tasks.

In the STM task, participants had to indicate whether a probe stimulus belonged to the encoded set or not. Chance level was 50%. A two-way rmANOVA was conducted with factors *session*, *level*, and their interaction. There was no effect of session ($F_{(2, 34)} = 0.45$, $p = .636$, $\eta^2 = .002$), a significant effect of level ($F_{(2, 34)} = 275.68$, $p < .001$, $\eta^2 = .717$), and no interaction ($F_{(4, 68)} = 0.43$, $p = .717$, $\eta^2 = .001$). Performance is plotted in Figure 7A. Overall, there was no change in short-term memory performance with sleep deprivation. The STM task was modelled after Habeck et al. (2004), for which a similar task was performed with fMRI following 48 h sleep deprivation. Their task used letters instead of symbols, and recognition accuracy had a significant effect of sleep deprivation, no effect of level, and no interaction. Comparing high work-load trials at BL to SD, Habeck et al. found increased BOLD activations in the anterior cingulate cortex (as well as thalamus and basal ganglia), whereas they found de-activations in parietal, temporal, and occipital regions. Our STM task used symbols, similar to Maurer et al. (2015). We had comparable accuracies for medium and low memory loads, however we were unable to replicate their correlation between log-transformed AFz theta increases from L1 to L3, and behavioral decreases in accuracy ($r = -0.08$, $p = .738$).

In the PVT, participants had to press a button whenever a counter started, every 2-10 s. The standard outcome measures for the PVT are reaction times (RT) and number of lapses (RTs > 0.5 s). Both RTs and lapses significantly increased with increasing sleep deprivation (Figure 7B), in line with previous studies. RTs increased from 295 ± 35 ms at BL to 359 ± 81 ms at SD ($t = 3.46$, $df = 16$, $p = .003$, $g = .82$). Reaction times were not corrected for recording system delays.

In the LAT, participants had to press a button as soon as they noticed the appearance of a gray circle in the attended visual hemifield. If their response was within 0.5 s, this counted as a correct trial; if a response was given up to 1 s after this, it was a late response; if no response was given, it was considered a lapse. Paired t-tests were conducted between all sessions. Mean RTs significantly increased with increasing sleep deprivation, from 372 ± 36 ms at BL to

408 ± 49 ms at SD (t = 4.16, df = 17, p = .001, g = 0.80). Reaction times were slower in the LAT compared to the PVT by 28% at BL, 28.5% at SR, and 20% at SD. The percentage of correct trials significantly decreased from BL to SR, and from BL to SD, but not between SR and SD, and likewise percentage of lapses increased (Figure 7C). Overall, LAT performance was negatively affected by sleep deprivation.

In the Speech Fluency Task, participants were presented 20 tongue twisters, and had to speak out loud each one as many times as possible within 10 s. Performance was measured by the number of correctly spoken words per second, and the number of mistaken words per second. The number of correct words significantly increased between BL and SR, and BL and SD. The number of mistakes significantly decreased between BL and SD, and SR and SD (Figure 7D). Performance therefore improved over the course of the experiment, likely due to a learning effect. Future studies should use new sentences for each session.

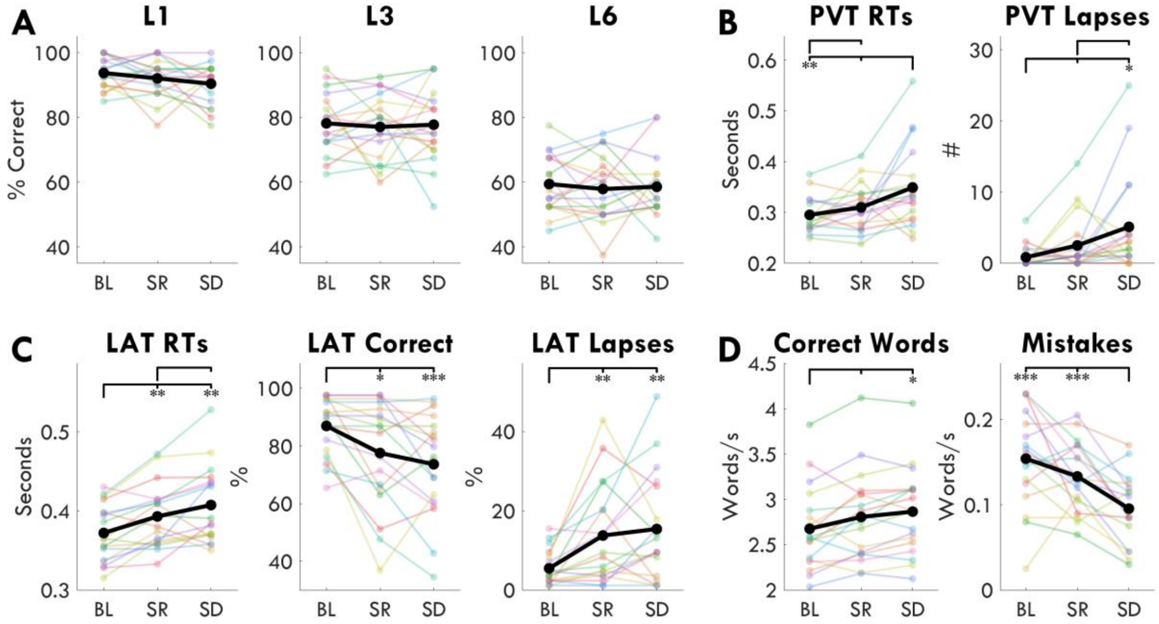


***Figure 7:*** ***A**: STM performance for every memory load level (1, 3, 6) at every session. The y-axis indicates percentage of correctly identified probes (both true positives and correct rejections). Thin lines indicate individual participants, the thick line indicates the grand mean. Chance level was 50%. No level showed a significant change from baseline. **B**: PVT performance. Left: mean RTs. Right: number of trials for which the RT > 0.5 s. **C**: LAT performance. Left: mean RTs in seconds. Middle: percentage of trials for which the RT was between 0.1 s and 0.5 s (i.e., while the stimulus was still visible). Right: percentage of trials for which no response was given. **D**: Speech Fluency Task performance. Left: rate of correct words per second across sessions. Right: rate of mistaken words per second across sessions. Asterisks indicate significant differences from paired t-tests between sessions, FDR corrected, such that: * p-value < .05, ** p-value < .01, *** p-value < .001. Acronyms: STM (short term memory task), LAT (lateralized attention task), PVT (psychomotor vigilance task), BL (baseline), SR (sleep restriction), SD (sleep deprivation), FDR (false discovery rate), RT (reaction times).*

### METHODS

#### Screening criteria

Applicants were screened prior to participating in order to: A) have a uniform, neurotypical population; B) avoid potential drop-outs due to adverse reactions to the experimental conditions; C) ensure participants' lifestyles were similar enough to the requirements of the control week (the week prior to each recording session) so as not to cause major disruptions; D) avoid any health or life conditions that could interact negatively with sleep deprivation or other experimental conditions; E) ensure participants were at least somewhat vulnerable to sleep deprivation in order to avoid floor effects.

Participants had to be between 18 and 25 (A). They had to be good sleepers with a PSQI ≤ 5 (Pittsburgh Sleep Quality Index (Buysse et al., 1989)), they had to have few night-time awakenings, and be resistant to adverse environmental conditions such as background noise or dim lights (B). They had to have a fairly regular sleep-wake rhythm, with an MCTQ score between 2 and 6.5 (Munich Chronotype Questionnaire (Roenneberg et al., 2015)), average sleep duration between 6 and 11 h, a preferred bedtime between 21:00-01:00 and wakeup time between 06:00-11:00 (C). Likewise, applicants were excluded if they regularly took naps (C). Applicants were excluded if they suffered any sleep-related disturbances or disorders such as insomnia or daytime sleepiness (D). They were excluded if they were pregnant or currently experiencing a difficult period in their life (stress, loss, etc.) (D). Applicants were excluded if they had any medical, psychological, or psychiatric conditions (B, D). Applicants had to have a healthy body weight, with a BMI (body mass index) between 18 and 30 (A, D). They were excluded if they had any physical impairment at the time of recording or had recently worn a long-term cast/bandage (D). They were further excluded if they had sensitive skin and might not tolerate wearing the EEG cap for extended periods of time (B). Applicants were excluded if they were currently or recently taking prescription medication, excluding contraceptives (A, D). They were excluded if they were regular recreational drug users, took prescription stimulants, if they were heavy consumers of alcohol (either daily consumption or occasional binge drinking), or if they were smokers (A, C). Applicants who consumed more than the equivalent of 3 cups of coffee per day were excluded (C). Applicants were excluded if they had prior experience with shift work, regular experience with changing time zones, or spending > 20 h awake (E). They were further excluded if they considered themselves resilient to sleep deprivation (E).

#### Questionnaires

A custom-built online survey tool, the Experiment Web Organizer for Questionnaires (EWOQ), was created for collecting questionnaire data through a web browser, written in React/typescript and hosted on Netlify and Google Cloud Platform. During the laboratory experiments, all questionnaires were filled out on a tablet, whereas the screening questionnaire and daily sleep reports were filled out on the participants' personal devices.

**Screening questionnaire**: The screening questionnaire was composed of 5 parts. First, applicants filled out a *Sensitive screening* questionnaire asking about medical history and drug use with yes/no questions. This data was not saved. If they passed, they continued to the *Pittsburgh Sleep Quality Index* (PSQI, (Buysse et al., 1989)) and then the *Munich Chronotype Questionnaire* (MCTQ, (Roenneberg et al., 2015)) to determine scores of regular sleep quality and individual chronotype, respectively. Then they filled out a more detailed questionnaire about their sleep quality (e.g. sensitivity to noise, night-time awakenings), and finally a questionnaire about their wake habits (e.g. caffeine consumption, time zone travel). After completing all questionnaires their data was saved and they were informed if they passed.

**Sleep reports**: Participants were asked to report every morning for the week prior to and including experiment nights: their sleep quality, whether they had any dreams, and what they had done the day prior. Data was not included in this paper.

**Task battery questionnaire**: Participants were asked after performing every task what their experience was during the task. Answers were given on a ~10 cm continuous slider with labels (indicated on the y axis in Figure 1). This included subjective sleepiness using the Karolinska Sleepiness Scale (Åkerstedt & Gillberg, 1990).

N.B.: only the PSQI, MCTQ and KSS are external, validated questionnaires. All others were created for this experiment and have not been tested on a broader population.

#### Recording equipment

Tasks were performed on a Lenovo ThinkPad P53 laptop (15.6" FHD, Intel Core i7-9750H) with Windows 10. The computer was kept at 50% volume and 100% brightness for all tasks. The tasks were programmed in Python v3.6.5 using the PsychoPy v3.2.4 toolbox. The code is available on GitHub[1]. Digital triggers were sent from the task computer to the EEG recording system via USB. Responses for the PVT and LAT were recorded with the USB-connected MilliKey™ button box. The Game was played on a 10.1" Huawei MediaPad T5, running Android Oreo.

**Net setup:** First, Natus® Genuine Grass® gold disk electrodes were placed below the chin for EMG and over the mastoids for the sleep polysomnography. The skin was scrubbed with Weaver Nuprep® gel and cleaned with skin disinfectant. The electrodes were filled with spes medica SAC2 Electrode Cream and secured to the skin with medical tape. Then, the circumference of the head, distance between nasion and inion and distance between earlobes was measured to establish net size and identify the position for Cz (50% distance from opposite landmarks). For the first recording bout, a few hairs were cut from the root at the Cz location to allow identification of this spot for the next experiment bout. The net (adult small, medium, or large, based on the head circumference) was placed on the head, centered on Cz, tightened as much as was comfortable and padded with cotton if requested. The skin under the ground and reference electrodes was additionally scrubbed and cleaned. All electrodes were then filled with ECI Electro-Gel™. Impedances were set to be < 5 kOhm for ground, reference, and external electrodes, and < 25 kOhm for all other electrodes. Gel was regularly refreshed every 4-6 hours during the sleep deprivation bout, and in the morning after each night of sleep.

#### EEG Preprocessing

Wake EEG data was cleaned and preprocessed in MATLAB (R2019b) using custom scripts[2] and the EEGLAB toolbox (v2019.1). The pipeline was largely based on the recommendations

[1] STM: https://github.com/snipeso/match2sample; LAT: https://github.com/snipeso/LAT; PVT: https://github.com/snipeso/pvt; Speech: https://github.com/snipeso/SFT;

[2] https://github.com/snipeso/Theta-SD-vs-WM

proposed by Makoto Miyakoshi[3] and standard practice in sleep research. The pipeline is represented in Figure 8.

First, data was low-pass filtered at 40 Hz using EEGLAB's default filter. A Kaiser notch filter was then applied to remove 50 Hz line noise and subsequent harmonics. Data was then down-sampled to 250 Hz. A 0.5 Hz high-pass Kaiser-window based FIR filter was then applied (0.25 Hz stopband, 60 dB stopband attenuation, 0.05 passband ripple).

The data was visually inspected to identify bad channels, bad time windows, and bad single-channel segments (i.e. snippets). Bad channels were considered as such that if they contained any non-physiological signals (anything not from the brain, muscles, or eyes) that occurred either continuously or repeatedly throughout the recording. Furthermore, external channels outside the EGI net were automatically removed (49, 56, 104, 113), as well as the face channels (126, 127). Bad time windows were any segments in time in which an artifact affected multiple channels at once, often due to body movements or brief muscle clenching. Bad snippets were non-physiological artifacts affecting only a few channels. Visual inspection was done by the same scorer (author SS) blinded to task, session, and participant. This manual preprocessing was chosen over automatic methods due to the unknown nature of sleep deprivation data, high variability across tasks in terms of data quality, and the possibility of true sleep intruding on wake.

Separately, the data was decomposed using independent component analysis (ICA; see *Independent component analysis* below), and artifact components were removed from the data after average referencing. Finally, removed channels and snippets were then interpolated such that 123 channels remained for each recording. Bad time windows were not included in the power analysis. Prior to the power analysis, 3 edge channels were removed (48, 119, 17), resulting in a final total of 120 channels. For source localization, the reference channel Cz was not included, as its exact coordinates were not available.

[3] https://sccn.ucsd.edu/wiki/Makoto's_preprocessing_pipeline retrieved December 2020.

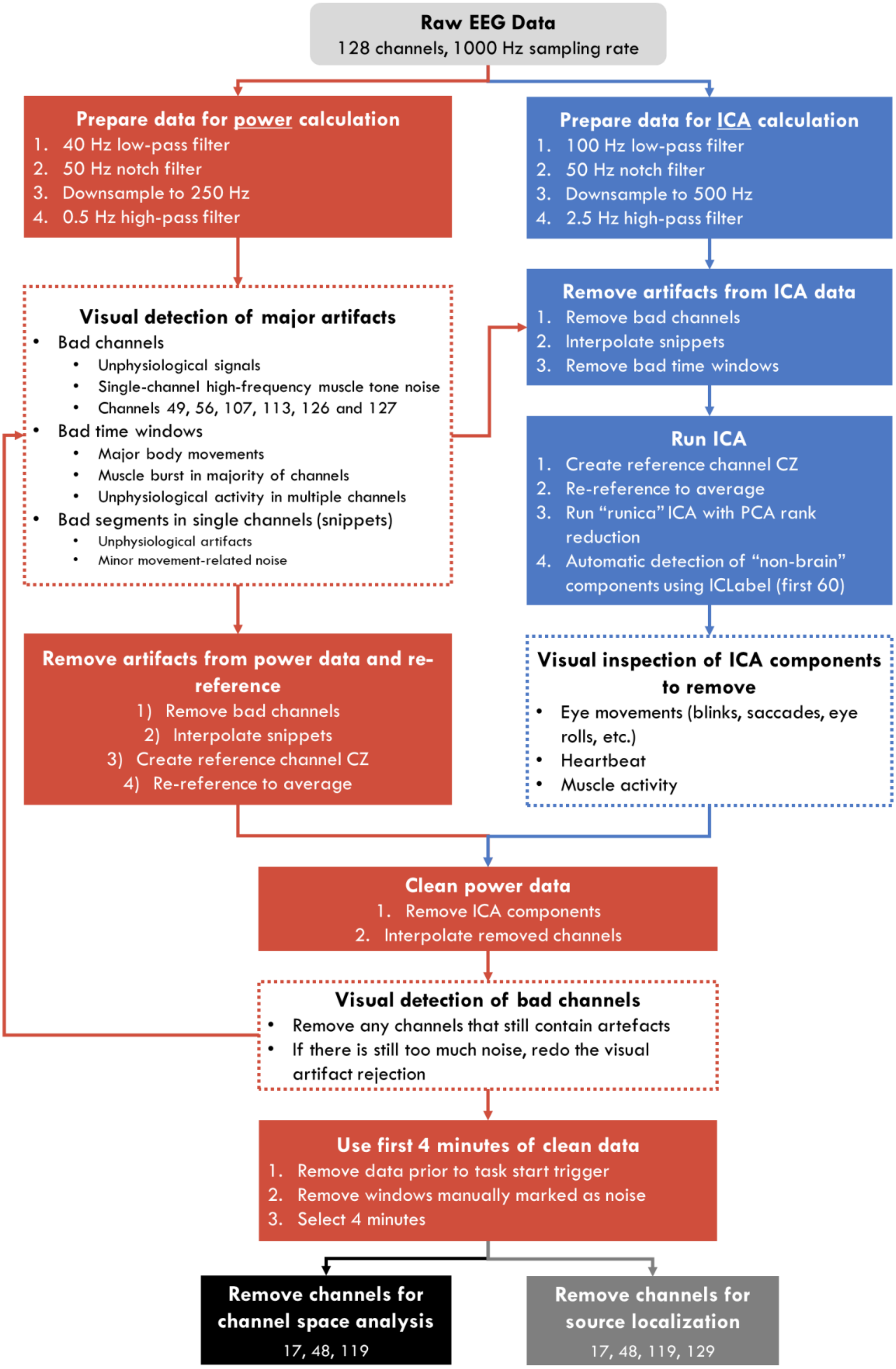


***Figure 8****: Pipeline for data preprocessing. Red steps are conducted on the data used for the power analysis. Blue steps are conducted on the data used for the ICA. White-filled steps involve manual work. Acronyms: EEG (electroencephalography), ICA (independent component analysis), PCA (principal component analysis).*

Overall, 4 ± 3 channels were removed on average per recording, out of 120. Figure 9 indicates the distribution per task and session. As expected, the Speech task had the highest number of channels removed. Unexpectedly, this difference became more attenuated with increasing sleep pressure.

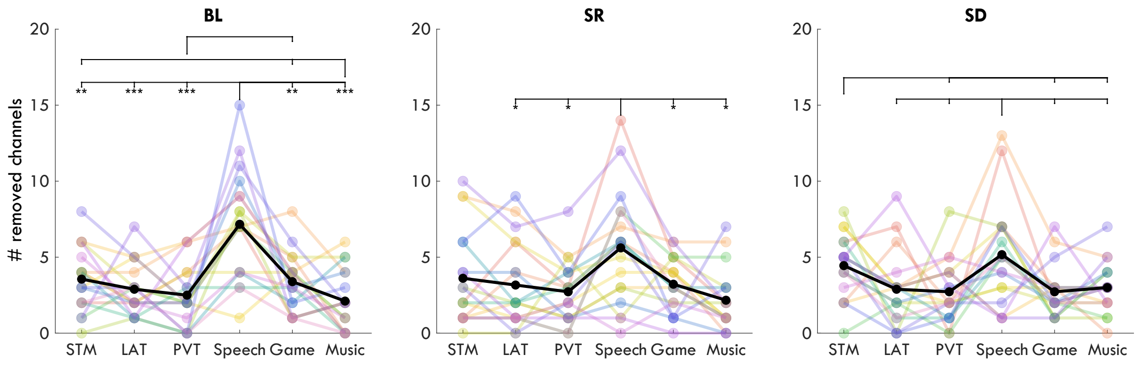


***Figure 9****: Number of channels removed per recording. Each colored line represents a participant. The black line is the group average. Asterisks indicate significant differences from paired t-tests between sessions, FDR corrected, such that: * p-value < .05, ** p-value < .01, *** p-value < .001. Acronyms: STM (short term memory task), LAT (lateralized attention task), PVT (psychomotor vigilance task), BL (baseline), SR (sleep restriction), SD (sleep deprivation), FDR (false discovery rate).*

#### Independent Component Analysis

Wake EEG data was cleaned from physiological artifacts using independent component analysis (ICA). This procedure allowed for the removal of eye movements (blinks, saccades, eye rolls), muscle activity (when limited to a few channels), heartbeat, and speech artifacts. For each recording, components were calculated based on data filtered between 2.5-100 Hz, and down-sampled to 500 Hz (Figure 8, blue), based on Dimigen (2020). Before ICA decomposition, 1) bad channels were removed, 2) bad snippets interpolated, 3) Cz restored, 4) bad time windows removed, and 5) all channels re-referenced to the average. EEGLAB's "runica" ICA algorithm was applied, with PCA (principal component analysis) rank reduction.

Using EEGLAB's ICLabel function (v1.2.4), the first 60 components were automatically classified as either brain data or artifacts. Components were marked for removal if they had a "brain" classification value lower than .1 but restored if they had a classification as "other" larger than .6 (i.e. an unknown component). Visual inspection (by SS, blinded to participant, session, and task) was then conducted to correct any misclassifications, or mark for removal additional bad components outside the first 60. These were not considered for automatic classification due to higher uncertainty for the classifier. After the components were removed, one final visual inspection of the data was done to remove any additional bad channels, and possibly repeat the preprocessing if notable artifacts remained in the data.

On average, 39 ± 12 components were removed from each recording (out of 106-122). Figure 10 indicates how many per participant, per session, per task. The Speech task had significantly more components removed (unsurprisingly), and the Music task the least. The majority of components removed were related to muscle artifacts.

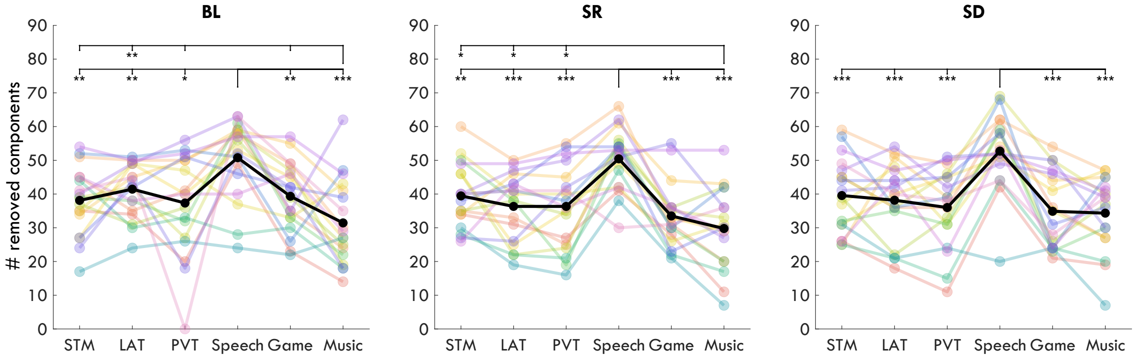


***Figure 10:*** *Number of removed components after ICA. Each color represents a single participant. Acronyms: ICA (independent component analysis), STM (short term memory task), LAT (Lateralized attention task), PVT (psychomotor vigilance task), BL (baseline), SR (sleep restriction), SD (sleep deprivation).*

#### EEG Analysis

The overall power analysis pipeline is provided in Figure 11, detailing the different steps undergone for each type of analysis.

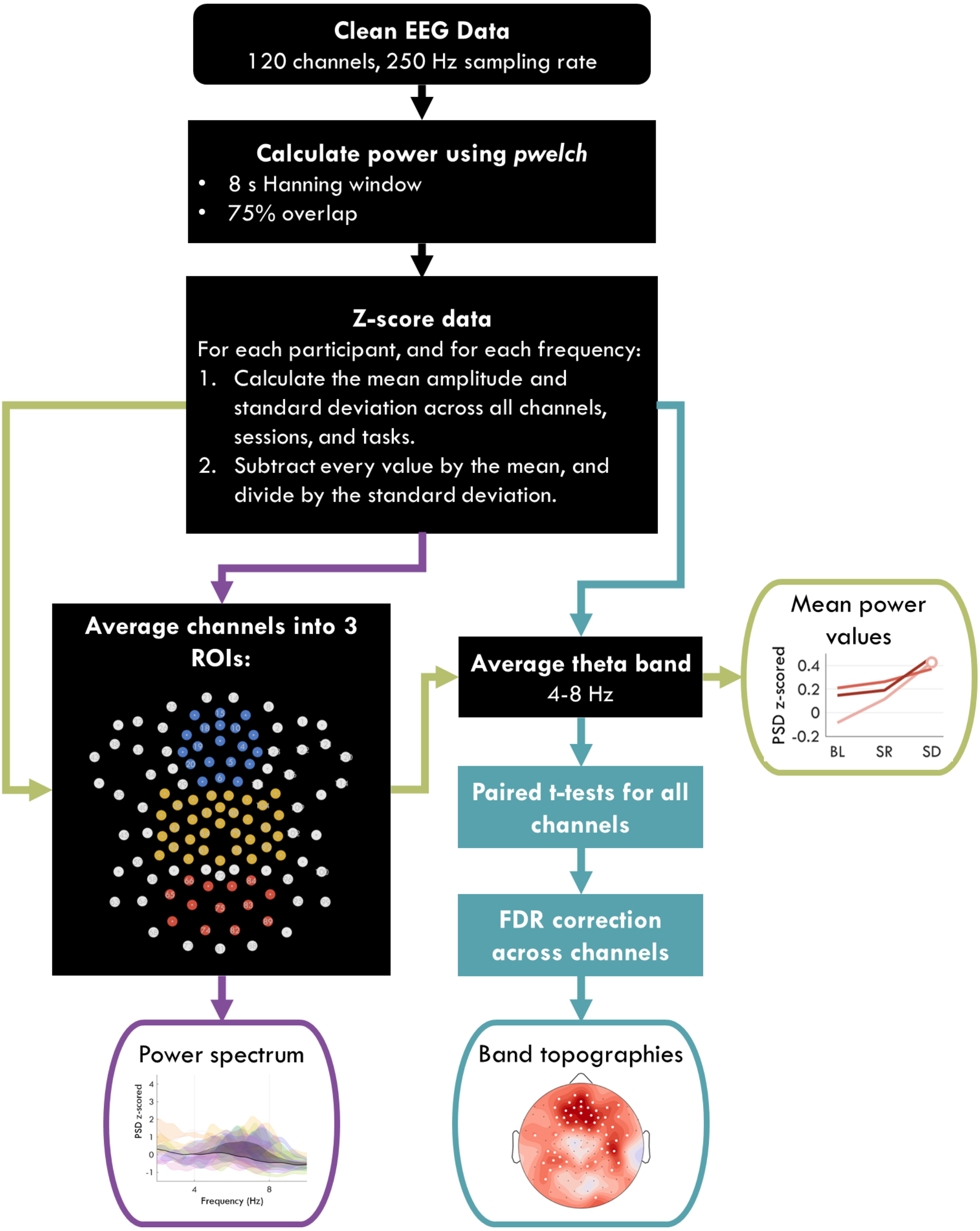


***Figure 11:*** *Pipeline for calculating power in the channel space. The paths marked by the colored arrows indicate the final steps for calculating the power spectrum, topographies, and ROI mean power values. For the ROIs, the blue dots indicate the front ROI, yellow the center, and red the back. Acronyms: ROI (regions of interest), FDR (false discovery rate), PSD (power source density), EEG (electroencephalography).*

**ROI channels:**

Front channels: 3, 4, 5, 6, 9, 10, 11, 12, 13, 15, 16, 18, 19, 20, 22, 23, 24, 112, 118, 124.

Center channels: 7, 30, 31, 35, 36, 37, 41, 42, 47, 51, 52, 53, 54, 55, 60, 61, 62, 78, 79, 80, 85, 86, 87, 92, 93, 97, 98, 103, 104, 105, 106, 110, 129.
Back channels: 65, 66, 69, 70, 71, 74, 75, 76, 82, 83, 84, 89, 90.

#### Z-Scoring

Z-scoring was done for the reasons outlined in the Results and Supplementary Results. The MATLAB code below outlines the exact steps involved:

```
% loop through participants
for indx_p = 1:tot_participants

    % loop through frequencies
    for indx_f = 1:tot_frequencies

        % retrieve for a given participant and given frequency
        % the power values of all sessions, tasks and channels
        DATA = data(indx_p, 1:tot_sessions, 1:tot_tasks, 1:tot_channels, indx_f);

        % calculate mean and standard deviation of all power values
        MEAN = mean(DATA, 'all');
        STD = std(DATA, 0, 'all');

        % calculate z-score
        z_data(indx_p, :, :, :, indx_f) = (DATA-MEAN)./STD;

    end
end
```

#### Source localization

The pipeline for source localization is depicted in Figure 12. To compute theta power for the source localization, we used a fast Fourier transform (FFT) with a Hanning taper, applied to each 8-s window.

To construct the forward model, we obtained a finite-element head model, implemented with the SimBio toolbox (Vorwerk et al., 2018), based on the segmentation of the template T1 MRI image from the Montreal Neurological Institute into gray matter, white matter, cerebrospinal fluid, scalp and skull. Subsequently, a standard 3D grid (10 mm spacing, 3294 voxels inside the head) and the head model were used to compute the leadfield matrix. To avoid depth bias, the leadfield was normalized.

As an inverse solution, we used the DICS beamformer technique, which enables the study of sources of oscillatory activation in the frequency domain (Gross et al., 2001). DICS is a linearly constrained minimum variance beamformer, which calculates the spatial filter using the sensor-level cross-spectral density (CSD) matrix, obtained by performing FFT at each leadfield matrix point. The spatial filter at each leadfield point is constructed in such a way that it passes activity from this location with unit gain, while attenuating activity from other locations (Gross et al., 2001). We first computed common spatial filters based on a cross-spectral density matrix obtained from pooled conditions, with a regularization parameter lambda set to 5%. By using common filter weights, we ensured that differences in source activity in contrasted conditions were not due to differences between condition-specific filters. The pre-computed common spatial filters were then applied independently to each condition. After projecting each recording to the source space, each frequency was z-scored for each participant (as done in the channel space).

**3D brain source maps**: To correct for multiple comparisons when performing t-statistics in the source space for visualization, we used a nonparametric clustering procedure (Maris and Oostenveld 2007; Maris 2012). This was done instead of FDR because it was only intended as a mask for the inflated brains plots, and not as hypothesis testing. First, independent samples t-tests for all voxels was done for the contrast of interest (2-tailed, $p < .05$). Next, significant neighboring voxels were clustered if they showed the same direction of effect. To assess the statistical significance of each cluster, a cluster-level test statistic was calculated by computing the sum of all t-values in the cluster. The significance of each cluster was estimated by comparing the cluster-level test statistic to a reference permutation distribution derived from the data. The reference distribution was obtained by randomly permuting the data 5000 times. The cluster *p*-value was estimated as the proportion of the elements in the reference distribution exceeding the cluster-level test statistic. For the visual representation of results, significant clusters of t-values were projected on the inflated brain surface. Due to uncertainty regarding the ability of surface EEG to detect deep brain structures' electrophysiological activity (thalamus, amygdala, etc.) these areas were covered in a patch and not included in the next analysis.

**Anatomical sources table**: To describe the source-level changes more easily, we additionally performed parcellation of the grid into 80 regions of interest (parcels), in accordance with the AAL atlas (Tzourio-Mazoyer et al. 2002). Regions of the basal ganglia and cerebellum were excluded from further analysis. The median power for each frequency across voxels was used for each anatomical area. Power for all theta frequencies was then averaged, and paired t-tests were conducted for each parcel, FDR correcting for multiple comparisons. Areas in Main Figure 6 only include those for which at least one contrast was significant.

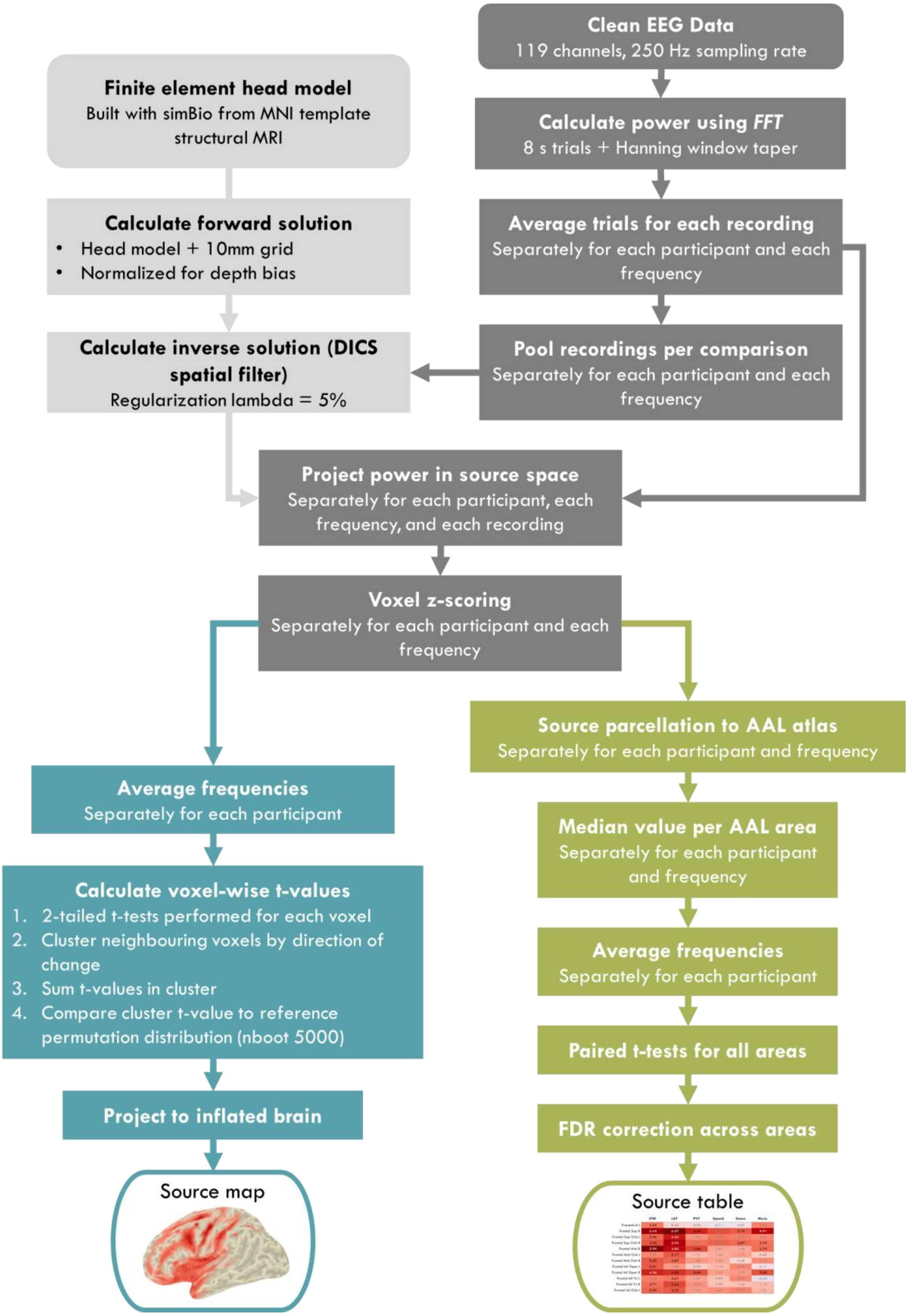


***Figure 12:*** *Pipeline for calculating change in theta power in the source space, both for plotting 3D brains and identifying source areas. Acronyms: FDR (false discovery rate), AAL (automated anatomical labelling atlas), DICS (dynamic imaging of coherent sources), MRI (magnetic resonance imaging), MNI (Montreal Neurological Institute), EEG (electroencephalography), FFT (fast Fourier transform).*

**The following areas were included prior to FDR correction:**

Precentral L, Precentral R, Frontal Sup L, Frontal Sup R, Frontal Sup Orb L, Frontal Sup Orb R, Frontal Mid L, Frontal Mid R, Frontal Mid Orb L, Frontal Mid Orb R, Frontal Inf Oper L, Frontal Inf Oper R, Frontal Inf Tri L, Frontal Inf Tri R, Frontal Inf Orb L, Frontal Inf Orb R, Rolandic Oper L, Rolandic Oper R, Supp Motor Area L, Supp Motor Area R, Olfactory L, Olfactory R, Frontal Sup Medial L, Frontal Sup Medial R, Frontal Med Orb L, Frontal Med Orb R, Rectus L, Rectus R, Insula L, Insula R, Cingulum Ant L, Cingulum Ant R, Cingulum Mid L, Cingulum Mid R, Cingulum Post L, Cingulum Post R, Hippocampus L, Hippocampus R, ParaHippocampal L, ParaHippocampal R, Calcarine L, Calcarine R, Cuneus L, Cuneus R, Lingual L, Lingual R, Occipital Sup L, Occipital Sup R, Occipital Mid L, Occipital Mid R, Occipital Inf L, Occipital Inf R, Fusiform L, Fusiform R, Postcentral L, Postcentral R, Parietal Sup L, Parietal Sup R, Parietal Inf L, Parietal Inf R, SupraMarginal L, SupraMarginal R, Angular L, Angular R, Precuneus L, Precuneus R, Paracentral Lobule L, Paracentral Lobule R, Heschl L, Heschl R, Temporal Sup L, Temporal Sup R, Temporal Pole Sup L, Temporal Pole Sup R, Temporal Mid L, Temporal Mid R, Temporal Pole Mid L, Temporal Pole Mid R, Temporal Inf L, Temporal Inf R.

#### Statistics

**ANOVAs**: all two-way repeated measures ANOVAs (analysis of variance) were calculated using MATLAB's Statistics and Machine Learning Toolbox. Greenhaus-Geisser corrected p-values were always used due to occasional violations of sphericity. Due to occasional missing data, not all analyses included every participant, therefore for each statistic the degrees of freedom (subscript in $F_{(A,\ B)}$) indicate the sample size ($N = \frac{B}{A} + 1$). No analysis had fewer than 16 participants, and the majority had all 18. Eta-squared ($\eta^2$) effect sizes were calculated using the Measures of Effect Size (MES) Toolbox[4] based on Hentschke & Stüttgen (2011).

**T-tests**: whenever only two conditions were being compared, paired t-tests were calculated to determine whether their difference was statistically significant. When more than one comparison is included in a plot (e.g. Main Figure 4A, 4D), FDR correction was applied. For channel-space plots, the FDR correction was done for the 120 channels of each plot. Hedge's g effect sizes are reported when t-values are described in the text. These were calculated using the MES toolbox.

**FDR correction**: Corrections for multiple comparisons was done by controlling for the *false discovery rate*, according to the procedure by Benjamini and Hochberg (1995). This was done using the Mass Univariate ERP Toolbox[5]. FDR was chosen over other methods because it required the fewest a-priori assumptions and thresholds (Groppe et al., 2011). All statistical tests were done with $\alpha = 5\%$.

---

[4] Harald Hentschke (2020). hhentschke/measures-of-effect-size-toolbox (https://github.com/hhentschke/measures-of-effect-size-toolbox), GitHub. Retrieved December 22, 2020.

[5] https://github.com/dmgroppe/Mass_Univariate_ERP_Toolbox
